# Parallel evolution under constraint shapes echinocandin resistance in *Candida auris*

**DOI:** 10.64898/2026.08.30.748140

**Authors:** Nicholas C. Cauldron, Erika N. Dort, Garrett M. Weeks, P. David Rogers, Christina A. Cuomo

**Affiliations:** Molecular Microbiology & Immunology, Brown University, Providence RI USA; Department of Pharmacy and Pharmaceutical Sciences, St. Jude Children’s Research Hospital, Memphis, TN; Infectious Disease and Microbiome Program, Broad Institute, Cambridge MA USA

## Abstract

Drug resistance emerges repeatedly in outbreaks of *Candida* fungal pathogens, but little is known about its origins or persistence. Here, we investigated the evolutionary processes shaping echinocandin resistance in *Candida auris*, a globally emerging and predominantly clonal fungal pathogen. Genome-wide association across over 600 isolates identified mutations in the β-1,3-glucan synthase gene *FKS1* as the most significant driver of resistance to an echinocandin drug. Ancestral reconstruction of this population traced shared resistance mutations among small groups typically consisting of 2-3 closely related isolates, but clusters could include up to 16 isolates. Nearly all resistant clusters consisted of isolates collected in the same year and region, consistent with local transmission. To further examine population-level selection, we measured adaptive signatures in *FKS1* and the highly diverged paralog *FKS2* across 22,000 genomes. This revealed excess nonsynonymous polymorphisms in *FKS1*, primarily due to independent, recurrent mutations at resistance hotspots, consistent with parallel evolution and incomplete fixation of adaptive alleles. In *FKS2*, there is no evidence of hotspots and little support for diversifying selection. Together, these results indicate that resistance mutations emerge under strong genetic constraint, with adaptation restricted to only one *FKS* homolog and predominantly at mutational hotspots.

**IMPORTANCE:** *Candida auris* is a critical public health threat due to its rapid global emergence and predisposition for multidrug resistance. While echinocandins are a first-line treatment, the emergence of resistance increasingly limits clinical options. Our study provides fundamental insights into the evolutionary processes driving this resistance. Through genomic analysis of thousands of clinical isolates, we demonstrate that high-level resistance is driven by mutation of the drug target, *FKS1*, and that this occurs through independent mutations at specific regions in the protein, termed hotspots. We show that while local transmission can expand resistant populations, drug adaptation occurs within an otherwise constrained landscape where amino acid changes are typically removed by selection. Furthermore, we establish that the paralog *FKS2* is functionally dispensable for resistance in *C. auris*, a difference from other *Candida* species. These findings define the patterns of resistance emergence for C. auris, offering a framework for genomic surveillance and mitigation strategies.

## INTRODUCTION

*Candida (Candidozyma) auris* is a recently emerged fungal pathogen distributed across all seven continents and can cause severe clinical complications (1–3). The World Health Organization lists *C. auris* as a critical priority pathogen due to these and other factors including acquisition of multidrug resistance (4).

Echinocandins are a cornerstone of treatments for invasive *C. auris* infection (5). Treatment failure often is caused by acquisition of resistance, most commonly arising from mutations in the drug target β-1,3-glucan synthase, encoded by *FKS* genes (6). Mutations most commonly emerge within one of three regions in the coding sequence (7, 8). These “hotspots” are embedded in the outer membrane and are situated adjacent to the glucan translocation channel (9–11). In *Saccharomcyes cerevisiae*, a distant yeast relative of *C. auris*, hotspot substitutions have been shown to confer resistance by both allosterically and sterically destabilizing echinocandin and glucan binding (12).

Multiple *FKS* paralogs are known to encode the same key enzyme (13), but it is not known why a subset of these paralogs are implicated in echinocandin resistance in certain *Candida* species or related yeasts (8, 14–16). The contributions of *FKS* paralogous genes to drug resistance are best delineated in *Candida glabrata* (taxonomically classified as *Nakaseomyces glabratus*), *S. cerevisiae,* and *Candida albicans*, which each possess three *FKS* genes. Resistance in clinical *C. glabrata* is linked to mutation of either *FKS1* or *FKS2* (17, 18), while resistance in *S. cerevisiae* is linked to *FKS2* mutations only when *FKS1* is deleted *in vitro* (19). Neither *FKS1* nor *FKS2* alone is essential in these species, but deletion of both genes is lethal to cells (13, 18). In *C. albicans*, only *FKS1* is essential and harbors the only known resistance mutations (20, 21), however deletion of *FKS2* or *FKS3 in vitro* is reported to reduce echinocandin susceptibility and moderately increase *FKS1* expression (22).

Similar to *C. albicans*, *C. auris* possesses two *FKS* genes, but only *FKS1* variants have been linked to drug resistance. While exposure to an echinocandin *in vitro* results in increased expression of *FKS2* but not *FKS1*, suggesting a possible role in drug response, transcript levels of *FKS2* are much lower than *FKS1* (23). However, the contributions of *FKS2* and its variants to echinocandin resistance in *C. auris* are not yet established.

Although hotspot mutations in *FKS1* are linked to *C. auris* echinocandin resistance, we know little of their relative persistence and contributions to outbreak clusters. The *C. auris* population structure is highly clonal, with outbreaks caused by three of the six lineages (24, 25). Since transmission of *C. auris* harboring *FKS1* mutations has been reported only rarely (26, 27), descriptions of echinocandin resistance frequency do not typically distinguish newly acquired mutations from those likely inherited from a related case (28–30). Thus, it is unknown how many independent origins have led to resistance and how stable resistance mutations are in the population. While most cases of echinocandin resistance are traced to mutations in *FKS1* hotspots, resistant isolates are occasionally reported without such mutations (24, 30, 31). Together, these indicate that our understanding of mechanisms driving the evolution of *C. auris* drug resistance is incomplete.

Here, we investigate the major mechanisms driving the emergence of echinocandin resistance in *C. auris*. Using genomic data and *in vitro* micafungin susceptibility as a surrogate for other echinocandins (14, 32), we demonstrate that *FKS1* is likely the sole driver of high-level *C. auris* echinocandin resistance. We show echinocandin resistance is associated predominantly with independent hotspot mutations in *FKS1*, but small clusters of likely regional transmission are inferred by ancestral reconstruction and supported by epidemiological data. By measuring selection among *C. auris* clades and comparison to selection on *FKS* homologs in related yeast pathogens, we find that clinically adaptive variants emerge in *FKS1* but not *FKS2*, and against a background of strong purifying selection. Since alternative resistance pathways are not detected in the conserved drug target paralog *FKS2*, and since recurrent mutations are highly localized in *FKS1*, we suggest echinocandin resistance evolves under strong constraint and follows highly predictable patterns which could be used as a proxy for detecting resistant isolates.

## RESULTS

### Identification of micafungin resistance mechanisms

To identify mechanisms of echinocandin resistance in *C. auris*, we performed a genome-wide association study using micafungin susceptibility data. We selected all public data that could be matched to available short-read, whole-genome sequencing data (Methods). This included a dataset of 681 isolates spanning the globe and including historic outbreaks (**Supplementary Fig. 1**). Micafungin resistance was detected based on the tentative minimum inhibitory concentration (MIC) breakpoint of >=4µg/mL (Methods) in only the three globally dominant lineages (Clades I, III, and IV).

We utilized this dataset to investigate how genetic variation associates with the level of micafungin resistance. To perform a genome-wide association study (GWAS), first, a “susceptibility index” was calculated for each isolate using its MIC (Methods; **Supplementary Table 1**). Then, non-synonymous variants identified from 669 single-sample isolates were tested for association with micafungin sensitivity. Rare variants (<0.05 allele frequency) were collapsed for each gene to test for a cumulative calculation (burden analysis, Methods).

This analysis identified a single major driver of resistance and secondary minor associations. While no single *FKS1* variant reached an allele frequency > 0.05, burden analysis revealed that variants in this gene were very strongly linked to micafungin resistance (**Table 1**). Outside of *FKS1*, the only significant hit after multiple test correction of p-values was the combined effect of rare variants in the putative asparaginase homolog *ASP1*. However, the effect size (beta) was negative, indicating variants may decrease micafungin resistance rather than increase (as in *FKS1*).

**Table 1.** Statistically significant GWAS variants from *in vitro* micafungin susceptibility of 669 or 617 (*FKS1* hotspot exclusive) *C. auris* isolates.

| Gene | Homolog | AF | Beta | Significance<br>* | Polymorphic<br>clades | Function |
| --- | --- | --- | --- | --- | --- | --- |
| **All isolates** |  |  |  |  |  |  |
| CJI82_02060 | *FKS1* | 0.13 | 2 | 7.62 x 10<sup>-49</sup> | I, III, IV | 1,3-β-glucan synthase |
| CJI82_01344 | *ASP1* | 0.01 | -1.9 | 0.019 | I, IV | Asparagine catabolism |
| ***FKS1* hotspot**<br>**exclusive** |  |  |  |  |  |  |
| CJI82_03261 | *OPT6* | 0.051 | 0.90 | 0.001 | I, III | Oligopeptide transporter |
| CJI82_05400 | *PBN1* | 0.045 | 1.21 | 0.003 | I, III | Catalysis in GPI anchor<br>processing |
| CJI82_02541 | *ECM27* | 0.043 | 1.19 | 0.010 | III | Regulation of calcium influx;<br>mutants papulacandin B<br>sensitive |
| CJI82_01962 | *MTR1* | 0.045 | 0.88 | 0.076 | I,III | Required for adhesion |
| CJI82_04140 | - | 0.045 | 1.22 | 0.090 | I | ATPase and ssDNA annealing |

\*Significance threshold < 0.1 by likelihood ratio test after Bonferroni multiple-test correction.

In our initial search, we identified seven isolates that were reported to be resistant to echinocandins yet lack an *FKS1* mutation. One isolate, upon re-testing, was found to be drug sensitive (B12779: 0.5 µg/mL). Genomic re-analysis of two other isolates with existing (B21247) or newly generated (B11784) data identified *FKS1* mutations associated with resistance (S639F and F635Y, respectively). Two of the other isolates are only reported as resistant to other echinocandins and not to micafungin (B11858: anidulafungin and caspofungin; B13905: caspofungin) (31). This re-analysis suggests that only two isolates may exhibit micafungin resistance and are lacking *FKS1* mutations: VPCI 550/P/14 (33) and VPCI 264/P/14 (B11218) (1).

To identify resistance genes independent of *FKS1* with an unbiased approach, we performed a stratified GWAS. We excluded any isolates harboring a mutation within a mutational hotspot of *FKS1* then repeated the correction for serial collections, yielding 623 isolates. While only two resistant isolates remained in the dataset based on the tentative breakpoint (**Supplementary Table 1**), we sought genes that may contribute to more intermediate sensitivities which may be on the path to higher-level resistance. Similar to the GWAS performed on all isolates agnostically of their *FKS1* allele, the most significant hits identified in the stratified GWAS are rare variant effects collapsed by gene (**Table 2**). Since evidence of parallel evolution is strongest at the gene-level (34, 35), strong GWAS candidates include genes with variants found in isolates with reduced micafungin susceptibility, for which multiple clades are represented. The relative significance of top hits in our GWAS was strongly affected by the minor allele frequency and the number of clades in which the gene is polymorphic. Genes significantly linked to reduced drug susceptibility have homologs in *S. cerevisiae* or *C. albicans* involved in oligopeptide transport (*OPT6*), GPI anchor processing (*PBN1*), or regulation of calcium influx (*ECM27*). With a loosened significance threshold, two additional hits are to an adhesin (*MTR1*) and an ATPase (*MGS1*). By controlling for the dominant effect of *FKS1* hotspot mutations, the stratified GWAS uncovered a set of candidate loci that may contribute to more subtle variation in micafungin susceptibility.

**Table 2.** Transmission clusters inferred by ancestry are consistent with available epidemiological data.

| Clade | Transmission cluster | Fks1 mutant | Isolates | Countries | SNPs apart | Weeks apart |
| --- | --- | --- | --- | --- | --- | --- |
| I | 1 | S639F | 2 | Kenya | 0 | 5 |
| I | 2 | S639F | 2 | UAE, Kenya | 0 | 20 |
| I | 3 | S639P | 3 | USA-New York | 0 | 0 - 1.3 |
| I | 4 | S639Y | 2 | USA-New York | 0 | 1.3 |
| I | 5 | S639Y | 2 | USA-New York | 0 | 2 |
| I | 6 | S639Y | 2 | USA-New York | 0 | 4.4 |
| III | 7 | F635dup | 2 | Kenya | 9 | 2 |
| IV | 8 | M1267I | 6 | Israel | 29 - 54 | 0 - 47.9 |
| IV | 9 | S639P | 2 | USA-Illinois | 57 | 62.9 - 112.1 |
| IV | 10 | S639P | 16 | Venezuela-Maracaibo | 15 - 47 | 0 - 163.3 |

### Recurrent mutation and local transmission of *FKS1*-linked micafungin resistance

While the relationship between *FKS1* mutation and echinocandin resistance is tightly associated, the genetic properties of its emergence are poorly described. To examine the evolutionary origins of resistance, we performed ancestral sequence reconstruction by tracing the types of mutations along a phylogeny of each clade estimated from whole-genome SNPs. Mutations mapped to internal branches are shared among descendant isolates (clusters), consistent with origin prior to their divergence (ancestral), whereas mutations on terminal branches are restricted to single isolates and are consistent with independent emergence (derived). Recurrence was quantified as the count of independent origins of each mutation.

The strong association of *FKS1* variants with resistance identified in our GWAS is supported by isolates from multiple clades exhibiting highly elevated MICs and an *FKS1* mutation. While mutations in *FKS1* were identified in five clades, the isolates with the highest resistance had mutations in hotspot 1. These mutations include multiple substitutions at a single amino acid (S639) associated with high resistance and a range of more moderate sensitivities associated with an amino acid duplication (F635dup) only present in clade III. There are a total of 58 isolates with a mutation in any of the three canonical mutational hotspots, all in clades I, III, and IV (**Fig. 1**). In these outbreak clades, all non-synonymous and non-singleton variants appear to be linked to elevated MIC measurements (>= 0.5 µg/mL) (**Fig. 1A**). In clades II and V, two non-synonymous *FKS1* mutations which are outside of the canonical hotspots and not linked to drug resistance are fixed. These were the only non-synonymous *FKS1* variants found in these clades, consistent with the absence of high-level resistance found therein (**Supplementary Fig. 1C**).

**Fig. 1.**
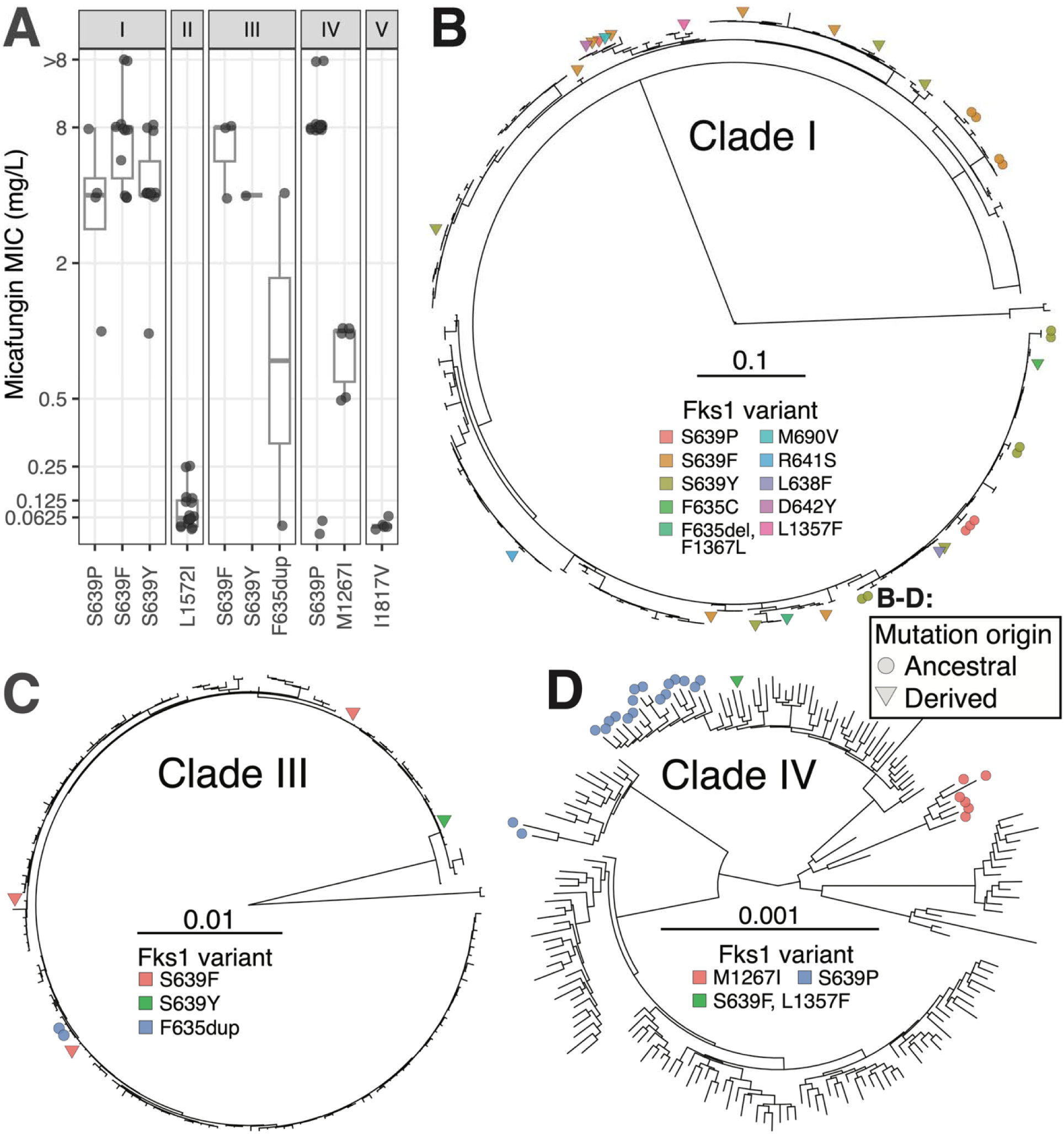
Resistance-associated Fks1 mutations originate both ancestral to case clusters and are derived independently in clinical *C. auris* populations. **A)** The micafungin MIC distribution of isolates harboring non-singleton Fks1 variants in each clade. **B–D)** Ancestral sequence reconstruction of each Fks1variant mapped onto maximum likelihood phylogenies for the three major clades was used to identify likely case clusters. Points at tips are colored by allele and shaped by inferred mutation origin: circles indicate mutations ancestral to a cluster, wherea triangles indicate derived mutations, independent to an isolate.

Overall, about half the total missense mutations in hotspots represent independent origins in those isolates (**Fig. 1B-D**, **Table 2**). Notably, one large cluster in Clade IV contains 16 phylogenetic siblings all harboring the S639P mutation, accounting for much of the imbalance of alleles and corresponding mutations for this site (**Fig. 1D**). This mutation is also the only one of ancestral origin, contributing to epidemiological clusters, in multiple clades (**Fig. 1B**, **Fig. 1D**). In contrast, the mutation that appears to have emerged the most recurrently was S639F, which is also the only variant found among isolates belonging to all three globally dominant clades (**Fig. 1**). A genotype with two mutations linked to resistance emerged only once, in one isolate (B12847: S639F and L1357F, clade IV), and neither mutation was found among closely related isolates.

To help distinguish whether isolate clusters descending from internal branches marked with mutations likely involved local transmission, the number of SNPs that differed between all isolate pairs was computed and was considered with available epidemiological data. Pairwise genetic distances were significantly different among the three clades (p < 2.2 x 10^-16^; Kruskal-Wallis test), primarily explained by elevated genetic distances between the clade IV isolates (**Supplementary Fig. 2**). So, the interpretation of genetic distance together with the collection date and region of each isolate is crucial to identify transmission clusters. Mutations found at internal branches, consistent with local transmission, include F635dup (1), S639P (3), S639Y (3), S639F (2), and M1267I (1) (**Supplementary Table 2**). Among these, descendants of all but one include isolates collected only from within the same region. Further, six such pairs and trios in Clade I were found to be 0 SNPs different and were collected within the same year. The two Clade III isolates harboring F635dup (**Fig. 1C**) were only nine SNPs different and were collected two weeks apart in Kenya.

The three candidate transmission clusters in Clade IV isolates appeared to be more distantly related. Although the six isolates with M1267I were all collected from Israel within a one-year period, including two pairs of isolates that were collected on the same date, they ranged 29-54 SNPs apart (**Table 2**). Similarly, a cluster of 16 isolates harboring S639P were all collected in Maracaibo, Venezuela and ranged 15-47 SNPs apart. Their collection dates cluster into two groups, with five collected in January – June 2015 and 11 collected in April – December 2012. The pair with 15 differences were collected only two weeks apart. While no pair of clade IV isolates was found with fewer than 15 SNP differences, the shared inheritance of *FKS1* alleles and longer time frames suggests the identification of transmission clusters.

### Diversity and selection on *FKS* genes

The β-1,3-glucan synthase gene family encodes two paralogs harboring the relevant enzyme domain in *C. auris*, *FKS1* and *FKS2*, and the role of *FKS2* is not well understood. While in *Candida glabrata*, echinocandin resistance is associated with mutations in both *FKS1* and *FKS2* genes, no *C. auris FKS2* variants are known to affect drug susceptibility. Across our dataset of 681 isolates with measured micafungin susceptibilities, we did not identify any non-synonymous or missense variants in *FKS2* that were associated with elevated MICs (>=0.5 µg/mL) **(Fig. 2A)**. However, since few isolates harbored mutations at this locus, sampling a larger population may reveal more rare evolutionary instances.

**Fig. 2.**
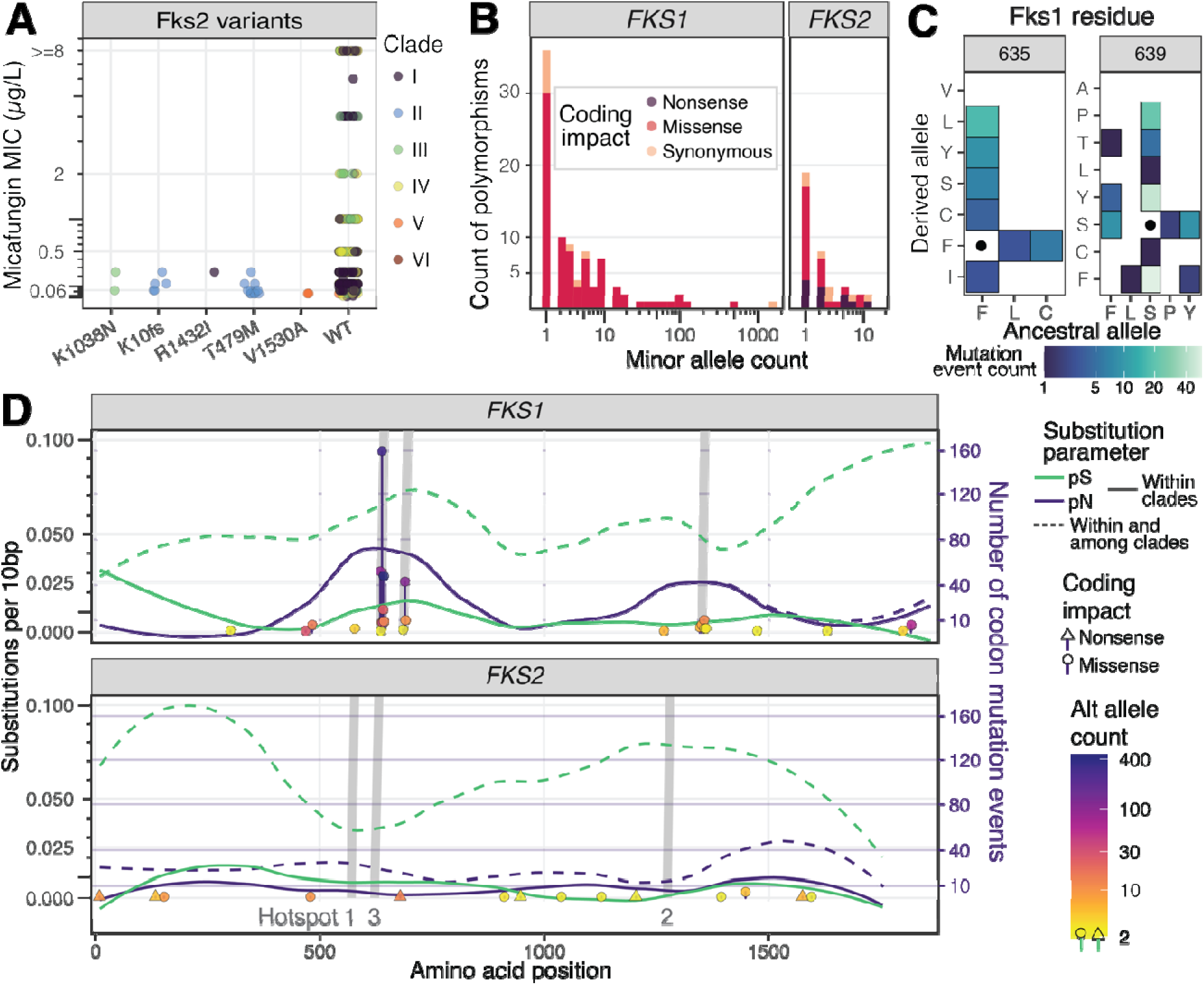
Estimates of diversity and selection in the global *C. auris* population differ at *FKS1* and *FKS2*. **A)** Micafungin MIC distribution of isolates harboring Fks2 mutations (n=669). **B)** Site frequency spectra for *FKS1* and *FKS2* (n=22,388). Bars are stacked and colored according to coding impact. **C)** Heatmap of recurrent mutation frequencies at the two most frequently mutated Fks1 residues, determined by ancestral sequence reconstruction (n=22,388). **D)** Substitution rate (horizontal lines; primary y-axis) and non-singleton SNPs (vertical lines; secondary y-axis indicating frequency of recurrence) in each paralog. Canonical hotspots in *FKS1* and their homologous coordinates in *FKS2* are highlighted in grey.

To more closely examine evolutionary pressure, we contrasted constraint on *FKS1* and *FKS2* across a wider global population of 22,388 isolates. We first compared allelic diversity and found that *FKS1* harbors greater diversity than *FKS2*. For *FKS1*, there are more isolates with variation in the population and sites in the alignment than in *FKS2* (**Fig. 2B**). Analyzing only non-synonymous mutations that are not fixed among clades, there were 30 sites with singleton alleles and 36 sites with non-singleton alleles in *FKS1*. In contrast, *FKS2* harbored only 17 or 15 such sites, respectively, despite similar estimates for numbers of possible non-synonymous sites across the two genes (*N*; **Table 3**).

**Table 3.** Selection pressure is stronger in *FKS1* than *FKS2* and targets mutational hotspots in the global *C. auris* population (n=22,389).

| minAC | *FKS1* (N=4574.3, S=1126.6) |  |  |  | *FKS2* (N=4215.7, S=1097.3) |  |  |  |
| --- | --- | --- | --- | --- | --- | --- | --- | --- |
|  | pN/pS | pN/pS non-HS+ | DoS | DoS non-HS+ | pN/pS | pN/pS non-HS+ | Dos | DoS non-HS+ |
| 1 | 1.4 | 0.86 | -0.37 | -0.19 | 0.86 | 0.82 | -0.31 | -0.31 |
| 2 | 2.1 | 0.83 | -0.28 | -0.04 | 0.47 | 0.47 | -0.26 | -0.27 |
| 3 | 1.8 | 0.66 | -0.24 | -0.01 | 0.33 | 0.33 | -0.24 | -0.25 |
| 13 | 2.7 | 0.74 | -0.23 | -0.02 | 0 | 0 | -0.22 | -0.23 |
| 14 | 2.7 | 0.74 | -0.23 | -0.02 | NA** | NA** | -0.23 | -0.24 |

\* Pn and Ps were calculated excluding polymorphisms fixed among clades
+ non-HS: the four columns with this label exclude canonical hotspot mutations in *FKS1/FKS2*
\*\* DoS could not be calculated because Pn = Ps = 0

Since recurrent mutations are consistent with adaptation, we contrasted amino acid space explored at frequently mutated sites in *FKS1* and *FKS2*. Ancestral sequence reconstruction revealed every amino acid residue in Fks2 mutated between the wild-type allele and only one alternate. But in Fks1, there could be multiple alternate alleles, each a single mutation in the codon (**Fig. 2C**). Fks1 residues 635 and 639 are mutated the most frequently, including reversions such as Y639S, and mutations among derived alleles such as Y39F. For Fks1-F635, residing in hotspot 1, six of the seven possible non-synonymous mutations within one mutational step are detected, and five of six are found at S639. Only F635V and S639A are not detected, each requiring an unfavorable transversion of thymine to guanine at the first codon position. At the homologous position in *S. cerevisiae*, F639V had a high selection coefficient, but S639A had the weakest selection coefficient among alternate alleles at that position (14). Since these mutations are possibly adaptive, their absence from the population is likely a result of limited accessibility compared to nucleotide transitions that may confer similar drug resistance.

We next determined whether positive or negative selection acts on *FKS1* and *FKS2*. We measured ongoing or transient selection by comparing the number of non-synonymous polymorphisms (Pn) across all clades and normalizing by synonymous (Ps) polymorphism counts which act as the baseline, neutral mutation rate. Since nonsynonymous sites are more abundant than synonymous, each is divided by the number of possible substitutions (N and S). When alleles that are not polymorphic within any clade are excluded from the analysis, only *FKS1* exhibits a signature of positive selection (pN/pS > 1) (**Table 3**). Considering multiple minimum allele frequencies at which substitutions are included, pN/pS ratios in *FKS1* range 1.4 - 2.7, while in *FKS2* they range 0 - 0.9. Therefore, our finding that *FKS1* but not *FKS2* is under positive selection in contemporary populations is robust across multiple genotyping quality thresholds.

Since *FKS1* variants conferring resistance tend to fall within canonical mutational hotspots, we searched for evidence of conserved hotspots in *FKS2*. Nonsynonymous substitutions are concentrated near hotspots *FKS1* but not in *FKS2*. Excluding the hotspots in *FKS1*, the signature of positive selection is attenuated toward neutrality, with pN/pS shifted downward to 0.7 - 0.9 (**Table 3**). There is virtually no shift when excluding homologous hotspot coordinates in *FKS2*, which harbor only one non-synonymous, and singleton, polymorphism (S574Y). The differences in selection near these hotspots is supported in deeper evolutionary time by the direction of selection statistic, in which omission of hotspots brought the estimated strength of selection to neutrality in *FKS1* but not in *FKS2*.

Since gene-wide estimates of selection may obscure regional effects, we used a windowed approach to scan for hotspots in *FKS1* and *FKS2* using pN/pS ratios and frequency of recurrent mutation following ancestral sequence reconstruction of translated sequences. In *FKS2*, we found no evidence of excess non-synonymous changes at known hotspot positions (**Fig. 2D**). We further analyzed variant positions and found a high number of variants in the hotspots of *FKS1* contrasted with no difference from the regional levels of variation around the homologous coordinates in *FKS2*. After local regression, the number of non-synonymous substitutions per non-synonymous site in *FKS1* ranges 0.3 - 0.5 in these regions (**Fig. 2D**). The sites most recurrently mutated are within hotspot 1, within which residues 635 and 639 are explained by minimums of 52 and 159 mutations, respectively. The minimum number of recurrent mutations in coding DNA sequences (CDS) were also determined as an agnostic way to identify mutational hotspots. In *FKS1*, we found recurrent mutations at 20 CDS sites, all of which are near a canonical hotspot (**Supplementary Table 3**). In contrast, there is only one position in *FKS2* with a recurrent mutation, and it is distant from coordinates homologous to the *FKS1* hotspots.

### Loss-of-function mutations in *FKS2*

Consistent with weaker selection acting on *FKS2*, we observed multiple independent, putative loss-of-function mutations in this gene. While no nonsense alleles were detected in *FKS1*, there were 11 in *FKS2,* resulting in a predicted loss of encoded function (**Fig 2B**). There were nine unique indels that shifted the coding frame, without any subsequent indels that restored the coding frame, present among 39 isolates. In addition, high impact SNPs that disrupted the coding frame were found in two isolates, one a mutation of the splice site and the second introduced a premature stop codon (**Supplementary Table 3**). Clades I – IV each harbored at least five isolates with a frame-shifting indel upstream of the enzyme domain (peptides 741 – 1511) suggesting the protein would be non-functional. Resistance data was only available for one of these alleles (K10fs); all isolates with this variant displayed very low micafungin sensitivities (<= 0.5 µg/mL) (**Fig 2A**). The independent losses of this gene suggest it is not strictly essential for some infections, while the inability to detect *FKS1* loss events suggests this gene is essential for pathogenicity.

To directly test the requirement for *FKS2* enzymatic activity, we disrupted the gene in a clade I background and measured *in vitro* drug susceptibility. We found no difference in micafungin susceptibility by broth microdilution and no observed change in MIC to micafungin or caspofungin by gradient diffusion strips (**Supplementary Fig. 3**). This data supports that *FKS2* appears dispensable for response to echinocandins in *C. auris*.

### *FKS* gene evolution among Saccharomycete relatives

Consistent with strong functional constraint, both *FKS1* and *FKS2* are dominated by purifying selection, with limited evidence for site-specific positive selection restricted to *FKS1*. To identify an evolutionary explanation for this contrast, we characterized the evolutionary history of *FKS* gene family members among human-pathogenic Saccharomycetes with reports of *FKS*-linked echinocandin resistance.

Phylogenetic analysis places *C. auris FKS2* as anciently diverged from *FKS1* and distantly related to *C. albicans FKS2* (**Fig. 3A**). The *C. glabrata* Fks2 is recently diverged and not the ortholog of these species. A whole-genome duplication in a common ancestor of *C. glabrata* and *S. cerevisiae* resulted in a second copy of *FKS1* named *FKS2* in both species; both genes are linked to clinical resistance. However, homologs of the gene named *FKS2* in the CTG clade of *Candida*, and in other lineages that diverged early from Saccharomycetes, have not been linked to clinical resistance. They appear to have originated from an independent gene duplication event in a common ancestor.

**Fig. 3.**
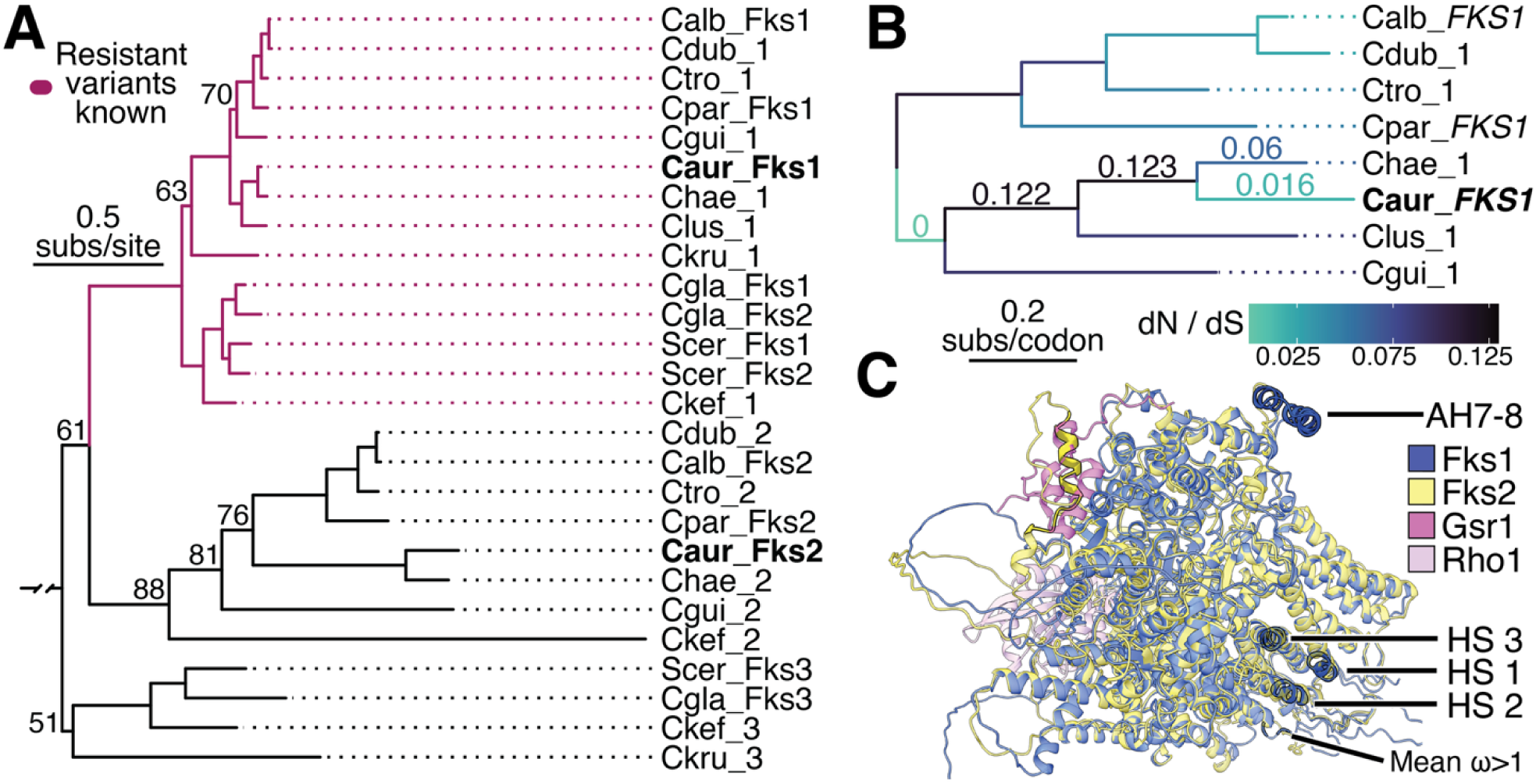
Orthologs of *C. auris* Fks1, but not its paralog Fks2, are linked to clinical echinocandin resistance. **A**) Phylogeny of Fks homologs in Saccharomycotina* with echinocandin resistance (bootstrap support >90 not shown; rooted on Scer* *GPI1*). Red branches indicate the paralog reportedly responsible. **B)** Nucleotide phylogeny from codon-aware alignment of *FKS1* including CTG clade species from A*. Branches are colored by the ratio of non-synonymous (dN) and synonymous (dS) substitutions and labeled leading to *C. auris FKS1*. **C)** Overlay of predicted *C. auris* Fks1 (blue) and Fks2 (yellow) protein structures. Depicted with binding locations of co-factors Gsr1 (9PE2; dark pink) and Rho1 (8WLA; light pink) predicted by homology. *Taxa in 1A and 1B abbreviated: *C. auris* (Cau), *C. lusitaniae* (Clus), *C. haemulonii* (Chae), *C. albicans* (Calb), *C. dubliniensis* (Cdub), *C. tropicalis* (Ctrop), *C. guillermondii* (Cgui), *C. parapsilosis* (Cpar), *C. krusei* (Ckru), *C. kefyr* (Ckef), *C. glabrata* (Cgla) and *S. cerevisiae* (Scer).

Consistent with these relationships, *C. auris FKS2* shares a more recent common ancestor with *C. albicans FKS2*, which is not linked to clinical drug resistance, than with *C. glabrata FKS2*. The genetic distance between *C. auris FKS1* and *FKS2* (2.1 substitutions per site) is much greater than the distance between *C. glabrata FKS1* and *FKS2* (0.1 s/s) or *FKS3* (1.6 s/s), a paralog for which no clinical resistance alleles are described.

To determine whether Fks homologs diverged under selection pressure, we calculated Dn/Ds within the sub-families of *FKS1* and *FKS2* genes (**Supplementary Table 4**). Reliable selection rates and codon trees for an *FKS1* alignment could be computed only within the CTG clade, a monophyletic group that is derived relative to *C. krusei* and *C. glabrata*. Reliable estimates for alignments of sub-families harboring *C. auris FKS2* and *C. glabrata FKS3*, and an alignment of the three *FKS* sub-families together, could not be computed because they were saturated for synonymous substitutions. Excluding *C. guillermondi* and *C. lusitaniae* improved but did not resolve saturation of the *FKS2* sub-family alignment.

In the CTG clade of *Candida*, *FKS1* homologs are under strong purifying selection (**Fig. 3B**). Allowing branch selection rates to be estimated freely was a stronger fit for the data than a uniform selection rate (**Supplementary Table 5**). But, in *FKS2*, the free-ratio model was not a significantly better fit than the one-ratio model. Performing this test in *FKS1* but excluding the same taxa as in *FKS2* still produced a significant result.

Although the signal of purifying selection was strong, an evolutionary model allowing positive selection at some sites in the *FKS1* alignment was a better fit than a model of uniform selection (**Supplementary Table 6**). We found that site-specific selection targeted the C-terminal: four sites with omega 1.47 – 1.49 were all within alignment sites 1849 – 1909 (**Supplementary Fig. 4**). These correspond to *C. auris FKS1* codons in the range of 1829-1882, residing in the extracellular component of the protein, well beyond any known resistance hotspot (nearest residues: 1351 – 1358) and all transmembrane helices (nearest residues: 1807-1827) in sequence space. In folded protein space, this region was predicted to reside at the end of the channel occupied by an extruding glucan (12), but outside of the distance to which it contacts any residues within the channel (**Fig. 3C**). The region harbors no substitutions among the approximately 22,000 genomes analyzed (nearest mutated residue: 1818).

To identify structural differences between *C. auris* Fks1 and Fks2, we modeled their structures using AlphaFold and constructed an overlay of the highest scoring models. The glycosyl transferase domain and mutational hotspots are strongly conserved, but there are two major differences within the accessory (AC) domain (**Fig. 3C**). The AC domain determines regulatory interactions, and in *S. cerevisiae* Fks1 (ScFks1) is resolved by Cryo-EM within residues 144-442 (36), which we found is homologous to *C. auris* Fks1 138-438. First, Fks1 harbors an additional pair of alpha helices in the region 242-272. In the catalytic conformation of the homologous ScFks1 structure, this region spans AH7-AH8, which are near the ‘latch loop’ that seals the active site during catalysis (12). This region is not fully resolved in any available cryo-EM structure of ScFks1, but it is consistent with the available AlphaFold-predicted structure (AF-A0A453J6Z3-F1-v6). Second, in all five predicted models of *C. auris* Fks2, an alpha helix (residues 72-87) appears to disrupt the binding pocket for Gsr1, a recently reported regulator of the enzyme complex in *S. cerevisiae* (37), either by obstruction (three models) or structural interaction (two models). No domains were predicted to disrupt the Gsr1 binding site in *C. auris* Fks1.

## DISCUSSION

Understanding the molecular basis and frequency of echinocandin resistance emergence is fundamental to predicting how it evolves. We investigated mechanisms of echinocandin resistance in clinical populations of *C. auris.* We show that i) *FKS1* is the clear driver of high-level micafungin resistance, ii) resistance commonly arises by recurrent mutations to hotspot sites, with contributions from regional transmission to expand resistant populations, and iii) adaptive mutations emerge, but do not rise to fixation, against a genetic background of strong purifying selection. Altogether, these results suggest that echinocandin resistance evolves primarily in the *FKS1* hotspot 1 within a highly constrained functional landscape.

A robust body of literature supports the role of *FKS* genes in echinocandin resistance among diverse fungi. In our data, we built upon prior work by doubling population size and performing a burden analysis, in which we identified *FKS1* as the main driver of high-level micafungin resistance in clinical *C. auris*. Of the two published GWAS on *C. auris* drug resistance, one recovered the association between *FKS1* and echinocandin resistance (38), but the other did not, likely due to paucity of isolates.

In our stratified GWAS removing the isolates with variants in mutational hotspots of *FKS1* linked to treatment failure (32), we identified several more novel gene candidates that may reduce micafungin susceptibility (38, 39). Putative roles include genes involved in stress response (40, 41) and cell wall synthesis or modification (42–44), as these pathways are closely linked to echinocandin resistance. In *C. albicans, OPT6* encodes an oligopeptide transporter that is induced two hours after phagocytosis, consistent with activation of a stress response pathway (45). In *S. cerevisiae*, *PBN1* is essential for the biosynthesis of GPI modifications, a pathway previously suggested as a promising target for antimicrobial drugs (46). *ECM27* may be involved in stress response pathways by interaction with Pdr5 (47), which exhibits increase sensitivity to echinocandins in *S. cerevisiae* and *C. albicans* when inhibited by FK506 (48–50), and deletion mutants are hypersensitive to an echinocandin (51). *MTR1* and CJI82_04140 have no close homologs in *S. cerevisiae*, but a *C. albicans* homolog of the former is required for adhesion (52).

We detected several candidate transmission events involving cases with an *FKS1* mutation and echinocandin resistance by combining ancestral state reconstruction, genetic distances, and epidemiological data (53, 54). Genetic distances among isolates involved in the candidate transmission events we detected are similar to prior reports of echinocandin resistant *C. auris* in Texas, Washington D.C., and California (26, 27). The high number of SNPs we found among clade IV isolates may have been influenced by the use of a clade I reference, as recent work using clade-specific reference genomes did not find this clade had elevated genetic distances relative to clades I or III (31), however, the larger collection time windows for these isolates might support their wider diversity.

Particularly in the presence of a strong selective pressure, Fks1 mutations that are inferred as ancestral could be independently acquired. However, since identical mutations rarely occur in a population at the same genomic position (55, 56), and since we have shown there are multiple utilized paths to resistance herein, we posit that accumulation of unlinked mutations drives divergence of clusters before *FKS1* mutation recurs. Therefore, an allele shared within a cluster of closely related isolates likely mutated only once. Consistent with this hypothesis, previous work found patient interaction histories were coherent with shared *FKS1* allele history and genetic distances (26). But there are likely some exceptions, as two pairs of closely related *C. auris* isolates with Fks1 mutations in New York did not share a recent healthcare history, suggesting recurrent mutation rather than transmission (57, 58).

While parallel evolution at the site-level is reported among clinical populations of fungi by *in vitro* evolution experiments (34, 59, 60), the frequency among clinical fungal populations is not well-described (35). We report extensive recurrent *FKS1* mutations in clinical populations of *C. auris*, consistent with parallel evolution. Our finding that most recurrent mutations are in hotspot 1 is consistent with prior population genomic analyses of *C. auris* indicating that it is the most frequently mutating hotspot (20, 30, 61), and that most mutations therein are under positive selection in *S. cerevisiae* (14). A study including diverse fungi reported that certain hotspot 1 positions were linked to resistance in 8 - 12 species (35). In *S. cerevisiae FKS1* hotspot 2, negative selection is predominant except for three sites where at least two unique amino acids accessible by one nucleotide mutation were under positive selection (14). Interestingly, those sites are homologous to the three sites in *C. auris FKS1* hotspot 2 in which we found recurrent mutations (1351, 1354, and 1357). Together, these data suggest that evolution of *FKS1* hotspot 2 is more highly constrained than hotspot 1.

Our analysis of recurrent mutations in *FKS1* identified several reversion mutations, indicating that some descendants of echinocandin resistant isolates reverted to the recent ancestral state of drug susceptibility. These reversions may occur to escape fitness penalties, which are commonly observed for drug resistance mutations in fungi (62). An alternative possibility is that the mutation was not fixed in the local population, and removal of the echinocandin selective pressure allowed resistance alleles to drift out of the population. However, since even this scenario is facilitated by relaxed selection, and given that reversions are statistically unlikely (63), these data are consistent with unknown fitness costs to maintaining echinocandin resistance (64). In *C. glabrata*, and *C. albicans, FKS* mutations are linked to growth defects and reduced virulence (65–67). In *C. auris*, two *FKS1* mutations have been shown to have some fitness tradeoffs, but no growth defects or reductions in virulence (68). We found no evidence of compensatory mutations, so it seems likely that a small fitness penalty results in the favoring of reversions (63, 64, 69, 70), however, the fitness effects of additional *C. auris FKS1* mutations would need to be directly measured to understand how any changes in selective pressure may facilitate reversions.

The *FKS* gene family paralogs exhibit different drug susceptibility phenotypes. A recently published Fks phylogeny indicated homologs cluster by species rather than by orthology (14), however, we show that the Fks2 lineage in the CTG clade is more deeply diverged than the recent paralogous divergence in *S. cerevisiae* and *C. glabrata.* While *FKS2* in *C. glabrata* and *C. albicans* are not orthologs, they both can affect echinocandin susceptibility (22). Our data establish that *C. auris* Fks2 appears to play no significant role in drug responses. This is consistent with recent reports that echinocandin response is reported to greatly vary between *C. auris* and *C. albicans*, likely attributed in part to differences in cell wall content (23, 71). Our models of folded *C. auris* Fks1 and Fks2 structures revealed few differences, suggesting that functional differences among homologs are driven by regulation. In *S. cerevisiae*, Fks1 synthesizes the majority of cell wall glucan in vegetative cells, while Fks2 and Fks3 collaborate to synthesize the majority of spore wall glucan (72, 73). In *C. albicans* and *C. auris* vegetative cells, Fks2 is expressed at much lower levels than Fks1 (2-3 fold and >32-fold, respectively) (21, 23), but further investigation including detection of mating may reveal a specific role.

Differences in the regulation of Fks homologs may be driven by regulatory sequences outside of the coding region or by structural differences influencing interactions among enzyme cofactors. In *S. cerevisiae*, enzyme cofactors Rho1 and Gsr1 bind to Fks1, modulating its expression (37). Rho1 activates all three Fks paralogs (ScFks1-3) (72), while Gsr1 has not yet been reported in complex with Fks3 (37). We identified two regions in which *C. auris* Fks2 differs from Fks1 that may impact the functions or binding of both cofactors. However, because the homologous residues in ScFks1 fall within unresolved disordered domains, it is difficult to infer how these structural differences contribute to differential regulation. Similarly, since in-depth evolutionary analysis of Gsr1 among pathogenic fungi has not yet been performed, authors caution that its regulatory role on Fks1 may be species-specific (12). Close investigation of the roles of enzyme cofactors in regulation of *C. auris* 1,3-β-glucan synthase activity is warranted.

In summary, genomic analyses reveal fundamental insights into the evolution of echinocandin resistance in *C. auris*. While echinocandin resistance emerges only rarely, a subset can be linked to transmission; the majority of cases are attributed to independent mutations at a small number of loci in *FKS1*, which is overall constrained under strong negative selection. This contrasts with the detected loss of function and experimental dispensability for resistance responses of the *C. auris FKS2* gene. Future experimental testing of regulatory networks involved in the echinocandin response may clarify the cellular roles of Fks homologs in *Candida* species, which will help develop strategies to mitigate the development of drug resistance in emerging fungal threats.

## METHODS

### Collection of whole-genome and micafungin susceptibility data

We performed a literature search to identify publications that both deposited genomic data in online repositories and included micafungin susceptibility test results. Drug susceptibility is measured by the minimum inhibitory concentration (MIC) of a drug needed to reduce growth by 50%. Data extracted from these published tables were supplemented with a dataset that included MIC measurements and sequencing data from the Centers for Disease Control (1, 30, 33, 58, 74–95). Sequencing data were generated by prior studies or were downloaded from NCBI using fasterq-dump from the SRA Toolkit v3.1.1 (96).

We resequenced two isolates (B11784 and B12779) reported to be micafungin-resistant despite harboring no *FKS1* mutations. We extracted high molecular weight DNA using a protocol modified from (97). Briefly, cells were treated R-Zymolyase (Zymo Research, California, USA) to digest the cell wall. DNA was purified with phenol and chloroform, and shipped to Plasmidsaurus (California, USA) for long- and short-read sequencing.

Since serially collected isolates from a single patient and strains from *in vitro* evolution experiments violate the statistical assumption of independence, we used publication details and notes provided by the CDC to select only one isolate from each case. We prioritized the isolate with the highest micafungin MIC and used the earliest collection date. To identify echinocandin resistance mechanisms driven by genes other than *FKS1*, a second subset of the population was also produced by first excluding isolates from the pool that harbored a mutation in any known *FKS1* hotspot and then re-applying the above selection criteria. These strategies resulted in collection of 681 unique isolates split into populations of 669 or 623 isolates agnostic of or excluding *FKS1*-hotspot mutants, respectively, that could be matched with micafungin MIC data.

### Collection of large-scale *C. auris FKS* genomic data

To obtain additional genomes for evolutionary study, we accessed *C. auris* isolates included in NCBI Pathogen Detection portal as of September 22, 2025. Each accession must meet several quality control filters to be included in species-wide clustering, resulting in a pool of high-quality genomes. If an assembled genome passes several more validation criteria, it is assigned to a cluster using k-mer distances followed by 50-SNP single-linkage clustering.

To reduce the download size, we used Magic-BLAST (98) to download only reads that aligned to either *FKS1* or *FKS2* in the B11205 reference genome (99). Using the NCBI BLAST+ suite v2.14.0+ (100, 101), we created a BLAST database that includes these genes and, to ensure adequate sequencing coverage at the 5’ and 3’ regions of each gene, the sequence 1kb upstream. We then processed these reads using the same workflow as for whole-genome data. After genotyping, only the whole genomes of isolates exhibiting *FKS* alleles that segregated within any clonal lineage were downloaded.

To determine which *C. auris* clade each NCBI SNP Cluster corresponded to, we searched our curated metadata for samples that were clustered and accessible in the NCBI Pathogen Detection Isolates Browser, and the parent SNP clusters were assigned a clade by majority rule. All isolates belonging to a SNP cluster with a determined clade then inherited this assignment. To determine the clade for isolates in SNP clusters that did not include any isolates with a previously annotated clade, we manually assigned it based on a phylogeny estimated from SNPs in *FKS1* and *FKS2* using FastTree (102) as described below. This tree was also used to visually assess cohesion of clade assignments. Isolates whose clade assignment conflicted between this phylogeny and the NCBI SNP cluster were discarded.

### Variant prediction

We mapped genomic reads to the B11205 reference genome and called variants using a previously published workflow (30, 99). Briefly, reads were mapped using BWA-mem v0.7.18 (103), processed with samtools v1.13 (104) to mark Illumina adapters and duplicates, and variants were then called with GATK v4.5.0.0 (105). We modified run parameters for GATK HaplotypeCaller according to suggestions provided by the GATK team for variant calling in fungal genomes, enabling --pileup-detection and --minimum-reads-per-alignment-start 75. Joint genotyping was performed using GenomicsDBImport and GenotypeGVCFs. The resulting file in Variant Call Format (VCF) was hard-filtered to omit low-quality variants using bcftools v1.22 for the following parameters: QD < 2.0, QUAL < 30.0, SOR > 3.0, FS (SNPs) > 60.0 or FS (indels) > 200, and MQ < 40.0 (30, 106). Since accurate phylogenies can be obtained from VCFs with fewer markers but higher stringency, we additionally censored genotypes that did not meet the following parameters: GQ < 50, alternate allele percentage < 0.8, and DP < 4. VCFs were annotated using SnpEff v5.2a (107) with a database built for the B11205 genome with the “Alternative_Yeast_Nuclear” codon table.

### Genome-wide association study

To perform genome-wide association, each VCF was converted to a variant matrix and then PLINK format using perl scripts as previously described (106), collapsing variants with a minor allele frequency < 0.05 by gene (vcfsToVariantMatrix.py and variantMatrixToPlink.pl; https://github.com/broadinstitute/broad-fungalgroup). To perform the association tests, we used GEMMA v0.98.5 (108, 109) with the parameters -lmm 4 -maf 0.01 -miss 1.0. These model parameters correct for underlying population structure using a kinship matrix in a multivariate linear mixed model.

We refer to transformed MIC values on which we performed the GWAS as a “susceptibility index” (**Table 1**). The minimum and maximum values of the susceptibility index were selected based on variation in the tested range of drug values among included studies. Some studies were included that determined micafungin MICs using 0.25 µg/mL as the minimum concentration, while others used 0.06 µg/mL. Similarly, some experimental designs allowed determination of the MIC at concentrations greater than 8.0 µg/mL, with a maximum value observed of 32.0 µg/mL, while in other designs any MIC larger than 8.0 µg/mL can only be reported as “>8.0” µg/mL. To preserve sample size while minimizing this source of error from experimental limitations, we transformed MICs below 0.25 µg/mL to a proportion of 1, and MICs greater than 8.0 µg/mL or reported as “>8.0” were both calculated as one dilution larger than 8.0 µg/mL. While the GWAS was also performed on raw MIC values, only results from the transformed values are shown since MICs are typically read within one dilution change of accuracy; the GWAS performed directly on MIC values is therefore more sensitive to error at higher concentrations.

### Phylogenetic analysis of *C. auris*

We estimated clade-specific phylogenies from concatenated SNP matrices. Samples were split into clade-specific VCFs using bcftools v1.23 (110). For each VCF, singleton variants were pruned and SNPs with less than 10% missing data were converted to FASTA format using vcfSnpsToFasta.py (https://github.com/broadinstitute/broad-fungalgroup) (24). Phylogenies were estimated under a generalized time-reversible model using FastTree v2.2.0 compiled with double precision (102). To estimate the reliability of each split in the tree, we used 1,000 resamples with the Shimodaira-Hasegawa test (111).

To perform ancestral sequence reconstruction, we used TreeTime v0.11.4 (112), which placed amino acid mutations of each translated gene sequence along clade phylogenetic trees according to the haplotype of each sample. *FKS1* haplotypes were extracted with samtools v1.20, bcftools v1.23, and translated with EMBOSS v6.6 (113). Since the ancestry of indels could not be reconstructed using the maximum likelihood approach in TreeTime, we manually inspected the clade-specific phylogenies near isolates harboring these variants (F635dup / L636FL) and assigned ancestral states by Fitch parsimony. The number of recurrent mutations was validated with RECUR v1.1.0 (114). For ancestral sequence reconstruction of the wider population of 22,388 genomes, only isolates with *FKS* alleles segregating within any clonal lineage were included since whole genomes were not downloaded for the entire population. To represent global diversity, the initial population of 669 isolates was also included.

Since the tools used to identify recurrent mutations are implemented for translated sequences, we used two other criteria to detect the minimum number of recurrent mutations in untranslated CDS. First, a genomic position with at least three alleles in a population must have mutated at least twice. Second, since sampled *C. auris* populations exhibit very limited evidence for recombination among clonal lineages since their divergence (25), positions where the same non-reference allele is identified among multiple clades are also consistent with recurrent mutation.

To detect selection, we contrasted rates of polymorphism within clades and divergence among clades in Python v3.11 and R v4.5.1 (115). We considered mutations identified in >= 95% of isolates within a subclade to be fixed (Dn or Ds), and variants with lower allele frequencies to be polymorphic (Pn or Ps). This tolerance allows for some missing genotype data and, rarely, isolates whose clade assigned by whole-genome SNP clustering is inconsistent with the Fks haplotypes expected for that clade. The ratio of nonsynonymous to synonymous polymorphism per site was calculated as (Pn/*N*)/(Ps/*S*), where *N* and *S* are the numbers of nonsynonymous and synonymous sites estimated with the unweighted Nei–Gojobori method (116) from the coding sequences of the reference genome (117). Scripts to calculate these statistics were generated, optimized, and debugged in part with Claude Code (Anthropic, Sonnet 4.6). The authors reviewed, tested, and take responsibility for the integrity and accuracy of the final code, which was deposited online: https://github.com/Neato-Nick/Cauris_FKS_evol. We calculated the direction of selection, which is the difference between the proportion of substitutions and polymorphisms that are nonsynonymous, using mkado v0.5.0 in vcf mode with --code-table 12 and *C. haemulonii* B16299 (SRR11092003) as the outgroup (118–121).

### *Candida auris* strains and growth conditions

*Candida auris* clade I strain 1c*FKS1*WT-11A (abbreviated 1cF), derived from the laboratory strain 1c (122), was used for all experiments. Relative to strain 1c, strain 1cF has the *FKS1*^S639F^ mutation corrected to wild-type sequence. Cultures were routinely grown in YPD (1% yeast extract, 2% peptone, 2% dextrose) at 35°C. For long-term storage, cultures were maintained in 25% glycerol at -80°C.

### Generation of *FKS2* deletion mutants

The *C. auris*-optimized episomal plasmid-induced Cas9 (EPIC) gene editing system (123) was used to create *C. auris FKS2* deletion mutants (plasmid pJMR19) via homology-directed repair.

#### Guide RNA design and plasmid preparation

A CRISPR-Cas9 guide RNA (gRNA) targeting the *FKS2* gene was designed in Benchling (124) using the Cand_auris_B8441_V2 genome for off-target analysis. The gRNA was ordered from Integrated DNA Technologies as two complementary, single-stranded 23-mer DNA oligos, each consisting of the 20-nucleotide gRNA sequence with SapI-compatible overhangs (**Supplementary Table 7**). The two oligos were duplexed and ligated into SapI (New England Biolabs)-digested pJMR19 plasmid (pJMR19-FKS2dis27) as previously described (125).

#### Repair template design and preparation

A gene disruption repair template was designed to integrate a stop codon cassette (5’-TAGNTAGNTAG-3’) 7 bp downstream of the *FKS2* start codon in the region adjacent to the Cas9 cut site as well as a silent blocking mutation to destroy the PAM site for the gRNA. The 662 bp repair template was ordered as a gBlock (Integrated DNA Technologies) and included 308 bp of sequence upstream and 330 bp of sequence downstream of the manipulation site (**Supplementary Table 6**). The gBlock repair template was PCR-amplified using the KAPA HiFi HotStart PCR Kit (Roche Diagnostics) with the manufacturer’s recommended cycling protocol and primers designed to amplify the gBlock (AMP primers; **Supplementary Table 6**). Eight PCR reactions were purified as previously described (122), and the reactions were pooled and concentrated using a standard ethanol precipitation protocol.

#### Transformations

Electroporation-based transformation of *C. auris* strain 1cF was performed as previously described (122) with two minor modifications. First, overnight *C. auris* cultures were grown to an OD_600_ of 2.2-2.8, and second, transformations were performed with two different ratios of plasmid to repair template (1µg:5µg and 0.5µg:4µg).

#### Transformant screening

Colony PCR was used to screen for positive transformants as described previously (122) using primers designed to amplify an 851 bp region in the *FKS2* gene surrounding the repair template integration site (CHECK primers; **Supplementary Table 6**). Purified PCR reactions were sent to Plasmidsaurus for amplicon sequencing using Oxford Nanopore Technology. Positive transformants were cured of the pJMR19 plasmid by culturing in 5 mL of YPD or RPMI-MOPS (0.2% glucose) at 35°C for up to six passages of 72 hours each.

### Antifungal susceptibility testing (AFST)

AFST was performed on the 1cF wild-type (WT) strain and two independent plasmid-cured mutant *FKS2* disruption strains (derived from different original transformant colonies) using a broth microdilution (BMD) approach for micafungin and a gradient diffusion strip approach for micafungin and caspofungin.

For AFST by BMD, micafungin (Sigma) was prepared in dimethyl sulfoxide (DMSO) following the Clinical and Laboratory Standards Institute M27 method modified as previously described (126, 127). An Infinite M Nano microplate reader (Tecan) was used to take optical density readings at 600 nm (OD_600_), and minimum inhibitory concentrations (MICs) were determined as the lowest concentration of micafungin at which there was 50% growth inhibition. Testing was performed in biological triplicate and technical duplicate. For each *C. auris* strain, the mean OD_600_ value from each drug concentration tested was calculated from the technical duplicates of each biological replicate and percent growth inhibition was calculated relative to the OD_600_ mean of the growth control (1% DMSO). The growth inhibition data was plotted as the biological replicate means ± standard deviation.

For AFST by gradient diffusion strips (Liofilchem), MICs were determined as previously described (128) and in biological triplicate.

### Selection of genomes from echinocandin-resistant Saccharomycotina

Taxa are referred to herein using their conventional, clinical names (e.g., *Candida glabrata*, *Candida auris, Candida krusei*) rather than recently revised genus assignments within Saccharomycotina (e.g., *Nakaseomyces*, *Candidozyma, Pichia*) (129). This is consistent with clinical and epidemiological literature in which “*Candida*” and traditional epithets are predominantly used (130, 131). Gene names were assigned according to the Candida Genome Database or the Saccharomyces Genome Database (132–134). Saccharomycetes with reported Fks1-related echinocandin resistance were identified based on reported echinocandin resistance or Fks1 variants (17, 30, 135–140).

Genomes were downloaded from FungiDB release 68 (141), or NCBI if a higher-quality assembly was available. To identify Fks homologues, the predicted proteomes were clustered using OrthoFinder v3.0.1b1 with the --c-homologs parameter to cluster genes into groups consisting of all homologs (142, 143). The parameter -S diamond_ultra_sens was also enabled, increasing sensitivity for the search step performed DIAMOND v.2.10 (144). We selected the hierarchical orthologous groups (HOGs) that harbored an Fks-named gene in either *C. auris* or *S. cerevisiae*, and were predicted to encode an enzymatic domain for glycosyltransferase family 48 (**Supplementary Table 4**) (145).

### Evolution among *FKS* homologs

Fks HOGs were aligned and analyzed together to determine phylogenetic distance among orthologs, and each ortholog separately for the analysis of selection. Protein alignments were performed with MAFFT v7.525 (146) and trimmed with ClipKit v2.11.4 (147), using default parameters for each. Codon alignments were generated by threading CDS sequences onto the original protein alignment using Pal2Nal v14 (148), then trimmed with the log file from ClipKit using a python script (https://github.com/jsharbrough/protTrim2CDS). Phylogenies were estimated using RAxML-ng v1.2.2 (149, 150) under the mutational model LG+I+G4, the best-fit model determined with ModelTest-NG v0.1.7 (151). Branch support was determined by estimation of up to 1000 bootstrap trees, stopping when Felsenstein bootstrap support values converged.

Branch-specific and site-specific selection was inferred using CODEML from PAML 4.10.10 (152). We used the parameters icode = 8 to enable the correct codon translation table, disabled the missing data filter, accounted for codon usage bias by setting CodonFreq = 2, set other parameters following published guidance (153), and deposited exemplar control files for branch and site tests in a public repository (https://github.com/Neato-Nick/Cauris_FKS_evol). Sites under positive selection were identified using the Bayes Empirical Bayes theorem for p(ω>1) ≥ 0.95 with model = 0 and NSsites = 8. All models were run in triplicate to assess convergence.

Folded protein structures for *C. auris* Fks1 and Fks2 were predicted with the AlphaFold 3 Server and visualized with ChimeraX (154, 155). Both proteins were aligned using ChimeraX matchmaker to an *S. cerevisiae* Fks1 structure solved by cryo-electron microscopy in complex with a short glucan chain (PDB: 9PE2) (12). An additional structure in complex with Rho1 (PDB: 8WLA) was also aligned for visualization (36).

## ACKNOWLEDGEMENTS

The authors thank Nancy A. Chow for providing drug susceptibility and genomic data of CDC isolates. The authors also thank Elizabeth Yee for assistance with DNA extractions and Paul Cao for assistance in the development and maintenance of computational workflows. This project has been funded in part with Federal funds from the National Institute of Allergy and Infectious Diseases, National Institutes of Health, Department of Health and Human Services, under awards R01-AI169066 to C.A.C and P. D. R, and F32-AI194727 to N. C. C.

## DATA AVAILABILITY

Comparative and population genomic analyses were performed using open-source software tools (Methods) or custom scripts that are publicly available (https://github.com/broadinstitute/broad-fungalgroup and https://github.com/Neato-Nick/Cauris_FKS_evol). This project used public sequence data from NCBI. Samples sequenced by this project, and additional samples from prior CDC sequencing publications, were deposited in the NCBI Sequence Read Archive and linked to relevant BioProject identifiers: PRJNA722434 (isolates B11784 and B12149), PRJNA638416 (isolate B17073), or PRJNA493622 (isolate B12779).

## REFERENCES

1. Lockhart SR, Etienne KA, Vallabhaneni S, Farooqi J, Chowdhary A, Govender NP, Colombo AL, Calvo B, Cuomo CA, Desjardins CA, Berkow EL, Castanheira M, Magobo RE, Jabeen K, Asghar RJ, Meis JF, Jackson B, Chiller T, Litvintseva AP. 2017. Simultaneous emergence of multidrug-resistant *Candida auris* on 3 continents confirmed by whole-genome sequencing and epidemiological analyses. Clin Infect Dis Off Publ Infect Dis Soc Am 64:134–140.

2. Chowdhary A, Jain K, Chauhan N. 2023. *Candida auris* genetics and emergence. Annu Rev Microbiol 77:583–602.

3. Rhijn N van, Gan E, Hepo-oja P, Wang X, Li J, Duggan S, Firer D, Alsharqi L, Gifford H, Steenwyk JL, Brackin AP, Abdolrasouli A, Borman AM, Cuomo CA, Fisher MC, Armstrong-James D, Farrer RA, Usher J, Rhodes J. 2026. Antarctic marine microplastics reveals environmental persistence and rapid evolution of *Candida auris*. bioRxiv 10.64898/2026.03.13.711634.

4. World Health Organization. 2022. WHO fungal priority pathogens list to guide research, development and public health action. 978-92-4-006024–1.

5. Pappas PG, Lionakis MS, Arendrup MC, Ostrosky-Zeichner L, Kullberg BJ. 2018. Invasive candidiasis. 1. Nat Rev Dis Primer 4:1–20.

6. Lee Y, Robbins N, Cowen LE. 2023. Molecular mechanisms governing antifungal drug resistance. Npj Antimicrob Resist 1:5.

7. Pfaller MA, Diekema DJ, Turnidge JD, Castanheira M, Jones RN. 2019. Twenty years of the SENTRY antifungal surveillance program: results for *Candida* species from 1997–2016. Open Forum Infect Dis 6:S79–S94.

8. Santana DJ, Cauldron NC, Rogers PD, Cuomo CA. 2026. Deciphering the multidrug resistance paradigm in *Candida auris*. Antimicrob Agents Chemother 70:e01062–24.

9. Hu X, Yang P, Chai C, Liu J, Sun H, Wu Y, Zhang M, Zhang M, Liu X, Yu H. 2023. Structural and mechanistic insights into fungal β-1,3-glucan synthase FKS1. 7955. Nature 616:190–198.

10. Zhao C-R, You Z-L, Chen D-D, Hang J, Wang Z-B, Ji M, Wang L-X, Zhao P, Qiao J, Yun C-H, Bai L. 2023. Structure of a fungal 1,3-β-glucan synthase. Sci Adv 9:eadh7820.

11. Li J, Liu H, Li J, Liu J, Dai X, Zhu A, Xiao Q, Qian W, Li H, Guo L, Yan C, Deng D, Luo Y, Wang X. 2025. Cryo-EM structure of the β-1,3-glucan synthase *FKS1*-Rho1 complex. Nat Commun 16:2054.

12. Ren Z, Chhetri A, Liu C, Offner S, Sharma K, Borgnia MJ, Im W, Yokoyama K, Lee S-Y. 2026. Structural basis of fungal β-1,3-glucan synthase inhibition by caspofungin. Nature 1–9.

13. Mazur P, Morin N, Baginsky W, el-Sherbeini M, Clemas JA, Nielsen JB, Foor F. 1995. Differential expression and function of two homologous subunits of yeast 1,3-β-D-glucan synthase. Mol Cell Biol 15:5671–5681.

14. Durand R, Torbey AG, Giguere M, Pageau A, Dubé AK, Lagüe P, Landry CR. 2026. Mutational landscape and molecular bases of echinocandin resistance in *Saccharomyces cerevisiae*. Genetics iyag055.

15. De Luca DG, Alexander DC, Dingle TC, Dufresne PJ, Fuller J, German GJ, Haldane D, Hoang LM, Jiao L, Kus JV, Li L, Malejczyk K, Sheitoyan-Pesant C, Stein M, Graham M, Van Domselaar G, Bharat A. 2026. Phylogenomic analysis and genetic mechanisms of antifungal resistance in clinical isolates of *Candida glabrata* (*Nakaseomyces glabratus*) from across Canada, 2013-2020. Microbiol Spectr 14:e0180325.

16. Yang F, Zhang L, Wakabayashi H, Myers J, Jiang Y, Cao Y, Jimenez-Ortigosa C, Perlin DS, Rustchenko E. 2017. Tolerance to caspofungin in *Candida albicans* is associated with at least three distinctive mechanisms that govern expression of *FKS* genes and cell wall remodeling. Antimicrob Agents Chemother 61:10.1128/aac.00071-17.

17. Garcia-Effron G, Lee S, Park S, Cleary JD, Perlin DS. 2009. Effect of *Candida glabrata FKS1* and *FKS2* mutations on echinocandin sensitivity and kinetics of 1,3-β-d-glucan synthase: implication for the existing susceptibility breakpoint. Antimicrob Agents Chemother 53:3690–3699.

18. Katiyar SK, Alastruey-Izquierdo A, Healey KR, Johnson ME, Perlin DS, Edlind TD. 2012. Fks1 and Fks2 are functionally redundant but differentially regulated in *Candida glabrata*: implications for echinocandin resistance. Antimicrob Agents Chemother 56:6304–6309.

19. Johnson ME, Katiyar SK, Edlind TD. 2011. New Fks hot spot for acquired echinocandin resistance in *Saccharomyces cerevisiae* and its contribution to intrinsic resistance of *Scedosporium* Species. Antimicrob Agents Chemother 55:3774–3781.

20. Park S, Kelly R, Kahn JN, Robles J, Hsu M-J, Register E, Li W, Vyas V, Fan H, Abruzzo G, Flattery A, Gill C, Chrebet G, Parent SA, Kurtz M, Teppler H, Douglas CM, Perlin DS. 2005. Specific substitutions in the echinocandin target Fks1p account for reduced susceptibility of rare laboratory and clinical *Candida* sp. isolates. Antimicrob Agents Chemother 49:3264–3273.

21. Garcia-Effron G, Park S, Perlin DS. 2009. Correlating echinocandin MIC and kinetic inhibition of *fks1* mutant glucan synthases for *Candida albicans*: Implications for interpretive breakpoints. Antimicrob Agents Chemother 53:112–122.

22. Suwunnakorn S, Wakabayashi H, Kordalewska M, Perlin DS, Rustchenko E. 2018. *FKS2* and *FKS3* genes of opportunistic human pathogen *Candida albicans* influence echinocandin susceptibility. Antimicrob Agents Chemother 62:10.1128/aac.02299-17.

23. Lara-Aguilar V, Rueda C, García-Barbazán I, Varona S, Monzón S, Jiménez P, Cuesta I, Zaballos Á, Zaragoza Ó. 2021. Adaptation of the emerging pathogenic yeast *Candida auris* to high caspofungin concentrations correlates with cell wall changes. Virulence 12:1400–1417.

24. Muñoz JF, Gade L, Chow NA, Loparev VN, Juieng P, Berkow EL, Farrer RA, Litvintseva AP, Cuomo CA. 2018. Genomic insights into multidrug-resistance, mating and virulence in *Candida auris* and related emerging species. 1. Nat Commun 9:5346.

25. Wang Y, Xu J. 2022. Population genomic analyses reveal evidence for limited recombination in the superbug *Candida auris* in nature. Comput Struct Biotechnol J 20:3030–3040.

26. Brown J, Le M, Tirol D, Kryger E, Villareal G, Fontela CQ, Torres A, Chiem T, Buchanan V, Kapsak CJ, Crumpler M, Zahn M. 2026. 546. Transmission of *Candida auris FKS1* mutations in Orange County, CA. Open Forum Infect Dis 13:ofaf695.019.

27. Lyman M, Forsberg K, Reuben J, Dang T, Free R, Seagle EE, Sexton DJ, Soda E, Jones H, Hawkins D, Anderson A, Bassett J, Lockhart SR, Merengwa E, Iyengar P, Jackson BR, Chiller T. 2021. Notes from the field: Transmission of pan-resistant and echinocandin-resistant *Candida auris* in health care facilities ― Texas and the District of Columbia, January–April 2021. MMWR Morb Mortal Wkly Rep 70:1022–1023.

28. Yadav A, Singh A, Wang Y, van Haren MH, Singh A, de Groot T, Meis JF, Xu J, Chowdhary A. 2021. Colonisation and transmission dynamics of *Candida auris* among chronic respiratory diseases patients hospitalised in a chest hospital, Delhi, India: A comparative analysis of whole genome sequencing and microsatellite typing. J Fungi 7:81.

29. Tian S, Wu Y, Li H, Rong C, Wu N, Chu Y, Jiang N, Zhang J, Shang H. 2024. Evolutionary accumulation of *FKS1* mutations from clinical echinocandin-resistant *Candida auris*. Emerg Microbes Infect 13:2377584.

30. Chow NA, Muñoz JF, Gade L, Berkow EL, Li X, Welsh RM, Forsberg K, Lockhart SR, Adam R, Alanio A, Alastruey-Izquierdo A, Althawadi S, Araúz AB, Ben-Ami R, Bharat A, Calvo B, Desnos-Ollivier M, Escandón P, Gardam D, Gunturu R, Heath CH, Kurzai O, Martin R, Litvintseva AP, Cuomo CA. 2020. Tracing the evolutionary history and global expansion of *Candida auris* using population genomic analyses. mBio 11:e03364–19.

31. Parnell LA, Dos Santos AR, Forsberg K, Lyman M, Misas E, Gade L, Sexton DJ, Litvintseva AP, Chow NA. 2026. Updated genomic epidemiologic description of *Candida (Candidozyma) auris*, United States. Emerg Infect Dis 32.

32. Perlin DS. 2015. Mechanisms of echinocandin antifungal drug resistance. Ann N Y Acad Sci 1354:1–11.

33. Sharma C, Kumar N, Pandey R, Meis JF, Chowdhary A. 2016. Whole genome sequencing of emerging multidrug resistant *Candida auris* isolates in India demonstrates low genetic variation. New Microbes New Infect 13:77–82.

34. Gerstein AC, Lo DS, Otto SP. 2012. Parallel genetic changes and nonparallel gene– environment interactions characterize the evolution of drug resistance in yeast. Genetics 192:241–252.

35. Bédard C, Pageau A, Fijarczyk A, Mendoza-Salido D, Alcañiz AJ, Després PC, Durand R, Plante S, Alexander EMM, Rouleau FD, Jordan DF, Jay A, Giguère M, Bernier M, Sharma J, Maroc L, Gervais NC, Menon ACT, Gagnon-Arsenault I, Bakker S, Rhodes J, Dufresne PJ, Bharat A, Sellam A, De Luca DG, Gerstein A, Shapiro RS, Quijada NM, Landry CR. 2025. FungAMR: a comprehensive database for investigating fungal mutations associated with antimicrobial resistance. Nat Microbiol 1–15.

36. Li J, Li J, Zhu A, Dai X, Liu J, Liu H, Xia Z, Dong Y, Qian W, Dai L, Guo L, Yan C, Deng D, Luo Y, Wang X. 2025. Structural-guided identification of two modulators of β−1,3-glucan synthase *FKS1*. Nat Commun 17:591.

37. Litsios A, Grys BT, Kraus OZ, Friesen H, Ross C, Masinas MPD, Forster DT, Couvillion MT, Timmermann S, Billmann M, Myers C, Johnsson N, Churchman LS, Boone C, Andrews BJ. 2024. Proteome-scale movements and compartment connectivity during the eukaryotic cell cycle. Cell 187:1490–1507.e21.

38. Wang Y, Xu J. 2024. Associations between genomic variants and antifungal susceptibilities in the archived global *Candida auris* population. 1. J Fungi 10:86.

39. Schikora-Tamarit MÀ, Gabaldón T. 2024. Recent gene selection and drug resistance underscore clinical adaptation across *Candida* species. 1. Nat Microbiol 9:284–307.

40. Satish S, Jiménez-Ortigosa C, Zhao Y, Lee MH, Dolgov E, Krüger T, Park S, Denning DW, Kniemeyer O, Brakhage AA, Perlin DS. 2019. Stress-induced changes in the lipid microenvironment of β-(1,3)-D-glucan synthase cause clinically important echinocandin resistance in *Aspergillus fumigatus*. mBio 10:10.1128/mbio.00779-19.

41. Shivarathri R, Jenull S, Stoiber A, Chauhan M, Mazumdar R, Singh A, Nogueira F, Kuchler K, Chowdhary A, Chauhan N. 2020. The two-component response regulator Ssk1 and the mitogen-activated protein kinase Hog1 control antifungal drug resistance and cell wall architecture of *Candida auris*. mSphere 5:10.1128/msphere.00973-20.

42. Zamith-Miranda D, Amatuzzi RF, Munhoz da Rocha IF, Martins ST, Lucena ACR, Vieira AZ, Trentin G, Almeida F, Rodrigues ML, Nakayasu ES, Nosanchuk JD, Alves LR. 2021. Transcriptional and translational landscape of *Candida auris* in response to caspofungin. Comput Struct Biotechnol J 19:5264–5277.

43. Jenull S, Shivarathri R, Tsymala I, Penninger P, Trinh P-C, Nogueira F, Chauhan M, Singh A, Petryshyn A, Stoiber A, Chowdhary A, Chauhan N, Kuchler K. 2022. Transcriptomics and phenotyping define genetic signatures associated with echinocandin resistance in *Candida auris*. mBio 13:e00799–22.

44. Dickwella Widanage MC, Singh K, Li J, Yarava JR, Scott FJ, Xu Y, Gow NAR, Mentink-Vigier F, Wang P, Lamoth F, Wang T. 2025. Distinct echinocandin responses of *Candida albicans* and *Candida auris* cell walls revealed by solid-state NMR. Nat Commun 16:6295.

45. Lorenz MC, Bender JA, Fink GR. 2004. Transcriptional response of *Candida albicans* upon internalization by macrophages. Eukaryot Cell 3:1076–1087.

46. Ashida H, Hong Y, Murakami Y, Shishioh N, Sugimoto N, Kim YU, Maeda Y, Kinoshita T. 2005. Mammalian PIG-X and yeast Pbn1p are the essential components of glycosylphosphatidylinositol-mannosyltransferase I. Mol Biol Cell 16:1439–1448.

47. Subba Rao G, Bachhawat AK, Gupta C. 2002. Two-hybrid-based analysis of protein–protein interactions of the yeast multidrug resistance protein, Pdr5p. Funct Integr Genomics 1:357–366.

48. Singh SD, Robbins N, Zaas AK, Schell WA, Perfect JR, Cowen LE. 2009. Hsp90 governs echinocandin resistance in the pathogenic yeast *Candida albicans* via calcineurin. PLOS Pathog 5:e1000532.

49. Tanabe K, Bonus M, Tomiyama S, Miyoshi K, Nagi M, Niimi K, Chindamporn A, Gohlke H, Schmitt L, Cannon RD, Niimi M, Lamping E. 2018. FK506 resistance of *Saccharomyces cerevisiae* Pdr5 and *Candida albicans* Cdr1 involves mutations in the transmembrane domains and extracellular loops. Antimicrob Agents Chemother 63:10.1128/aac.01146-18.

50. Iyer KR, Robbins N, Cowen LE. 2022. The role of Candida albicans stress response pathways in antifungal tolerance and resistance. iScience 25.

51. Lussier M, White A-M, Sheraton J, di Paolo T, Treadwell J, Southard SB, Horenstein CI, Chen-Weiner J, Ram AFJ, Kapteyn JC, Roemer TW, Vo DH, Bondoc DC, Hall J, Zhong WW, Sdicu A-M, Davies J, Klis FM, Robbins PW, Bussey H. 1997. Large scale identification of genes involved in cell surface biosynthesis and architecture in Saccharomyces cerevisiae. Genetics 147:435–450.

52. Desai JV, Bruno VM, Ganguly S, Stamper RJ, Mitchell KF, Solis N, Hill EM, Xu W, Filler SG, Andes DR, Fanning S, Lanni F, Mitchell AP. 2013. Regulatory role of glycerol in *Candida albicans* biofilm formation. mBio 4:10.1128/mbio.00637-12.

53. Lemieux JE, Siddle KJ, Shaw BM, Loreth C, Schaffner SF, Gladden-Young A, Adams G, Fink T, Tomkins-Tinch CH, Krasilnikova LA, DeRuff KC, Rudy M, Bauer MR, Lagerborg KA, Normandin E, Chapman SB, Reilly SK, Anahtar MN, Lin AE, Carter A, Myhrvold C, Kemball ME, Chaluvadi S, Cusick C, Flowers K, Neumann A, Cerrato F, Farhat M, Slater D, Harris JB, Branda JA, Hooper D, Gaeta JM, Baggett TP, O’Connell J, Gnirke A, Lieberman TD, Philippakis A, Burns M, Brown CM, Luban J, Ryan ET, Turbett SE, LaRocque RC, Hanage WP, Gallagher GR, Madoff LC, Smole S, Pierce VM, Rosenberg E, Sabeti PC, Park DJ, MacInnis BL. 2021. Phylogenetic analysis of SARS-CoV-2 in Boston highlights the impact of superspreading events. Science 371:eabe3261.

54. Gontjes KJ, Singh A, Sansom SE, Boyko JD, Smith SA, Lautenbach E, Snitkin E. 2025. Phylogenetic context of antibiotic resistance provides insights into the dynamics of resistance emergence and spread. J Infect Dis 232:e992–e1002.

55. Kimura M, Crow JF. 1964. The number of alleles that can be maintained in a finite population. Genetics 49:725–738.

56. Ma J, Ratan A, Raney BJ, Suh BB, Miller W, Haussler D. 2008. The infinite sites model of genome evolution. Proc Natl Acad Sci 105:14254–14261.

57. Ostrowsky B. 2020. *Candida auris* isolates resistant to three classes of antifungal medications — New York, 2019. MMWR Morb Mortal Wkly Rep 69.

58. O’Brien B, Liang J, Chaturvedi S, Jacobs JL, Chaturvedi V. 2020. Pan-resistant *Candida auris*: New York subcluster susceptible to antifungal combinations. Lancet Microbe 1:e193–e194.

59. Carolus H, Pierson S, Muñoz JF, Subotić A, Cruz RB, Cuomo CA, Van Dijck P. 2021. Genome-wide analysis of experimentally evolved *Candida auris* reveals multiple novel mechanisms of multidrug resistance. mBio 12:10.1128/mbio.03333-20.

60. Hirayama T, Miyazaki T, Sumiyoshi M, Ito Y, Ashizawa N, Takeda K, Iwanaga N, Takazono T, Yamamoto K, Izumikawa K, Yanagihara K, Makimura K, Tsukamoto K, Kohno S, Mukae H. 2023. Echinocandin resistance in *Candida auris* occurs in the murine gastrointestinal tract due to *FKS1* mutations. Antimicrob Agents Chemother 67:e01243–22.

61. Chowdhary A, Prakash A, Sharma C, Kordalewska M, Kumar A, Sarma S, Tarai B, Singh A, Upadhyaya G, Upadhyay S, Yadav P, Singh PK, Khillan V, Sachdeva N, Perlin DS, Meis JF. 2018. A multicentre study of antifungal susceptibility patterns among 350 *Candida auris* isolates (2009–17) in India: role of the ERG11 and FKS1 genes in azole and echinocandin resistance. J Antimicrob Chemother 73:891–899.

62. Hawkins NJ, Fraaije BA. 2018. Fitness penalties in the evolution of fungicide resistance. Annu Rev Phytopathol 56:339–360.

63. Teotónio H, Rose MR. 2001. Perspective: reverse evolution. Evolution 55:653–660.

64. Björkman J, Nagaev I, Berg OG, Hughes D, Andersson DI. 2000. Effects of environment on compensatory mutations to ameliorate costs of antibiotic resistance. Science 287:1479– 1482.

65. Ben-Ami R, Garcia-Effron G, Lewis RE, Gamarra S, Leventakos K, Perlin DS, Kontoyiannis DP. 2011. Fitness and virulence costs of *Candida albicans* FKS1 hot spot mutations associated with echinocandin resistance. J Infect Dis 204:626–635.

66. Singh-Babak SD, Babak T, Diezmann S, Hill JA, Xie JL, Chen Y-L, Poutanen SM, Rennie RP, Heitman J, Cowen LE. 2012. Global analysis of the evolution and mechanism of echinocandin resistance in *Candida glabrata*. PLOS Pathog 8:e1002718.

67. Arastehfar A, Daneshnia F, Hovhannisyan H, Cabrera N, Jusuf S, Salehi M, Mansour MK, Gabaldón T, Shor E, Perlin DS. 2026. Multidimensional assessment of in-host fitness costs of echinocandin resistance in the opportunistic fungal pathogen *Candida glabrata* reveals the niche-specific requirement for *FKS1* and *FKS2* during infection and gut colonization. Antimicrob Agents Chemother 0:e01801–25.

68. Ross ZK, Alsayegh S, Zhao Y, McPherson S, Munro CA, Lorenz A. 2025. *In vitro* evolution of caspofungin resistance in *Candidozyma auris* via *FKS1* hotspot I mutations results in moderate fitness trade-offs but no reduction in virulence. Microbiol Res 301:128322.

69. Schulz zur Wiesch P, Engelstädter J, Bonhoeffer S. 2010. Compensation of fitness costs and reversibility of antibiotic resistance mutations. Antimicrob Agents Chemother 54:2085–2095.

70. Pennings PS, Ogbunugafor CB, Hershberg R. 2022. Reversion is most likely under high mutation supply when compensatory mutations do not fully restore fitness costs. G3 GenesGenomesGenetics 12:jkac190.

71. Dickwella Widanage MC, Singh K, Li J, Yarava JR, Scott FJ, Xu Y, Gow NAR, Mentink-Vigier F, Wang P, Lamoth F, Wang T. 2025. Distinct echinocandin responses of *Candida albicans* and *Candida auris* cell walls revealed by solid-state NMR. Nat Commun 16:6295.

72. Ishihara S, Hirata A, Nogami S, Beauvais A, Latge J-P, Ohya Y. 2007. Homologous subunits of 1,3-beta-glucan synthase are important for spore wall assembly in *Saccharomyces cerevisiae*. Eukaryot Cell 6:143–156.

73. Lee-Soety JY, Resch G, Rimal A, Johnson ES, Benway J, Winter E. 2024. The MAPK homolog, Smk1, promotes assembly of the glucan layer of the spore wall in S. cerevisiae. Yeast 41:448–457.

74. Escandón P, Chow NA, Caceres DH, Gade L, Berkow EL, Armstrong P, Rivera S, Misas E, Duarte C, Moulton-Meissner H, Welsh RM, Parra C, Pescador LA, Villalobos N, Salcedo S, Berrio I, Varón C, Espinosa-Bode A, Lockhart SR, Jackson BR, Litvintseva AP, Beltran M, Chiller TM. 2019. Molecular epidemiology of *Candida auris* in Colombia reveals a highly related, countrywide colonization with regional patterns in amphotericin B resistance. Clin Infect Dis 68:15–21.

75. Chen X-F, Zhang H, Liu L-L, Guo L-N, Liu W-J, Liu Y-L, Li D-D, Zhao Y, Zhu R-Y, Li Y, Dai R-C, Yu S-Y, Li J, Wang T, Dou H-T, Xu Y-C. 2024. Genome-wide analysis of in vivo-evolved *Candida auris* reveals multidrug-resistance mechanisms. Mycopathologia 189:35.

76. Ben Abid F, Salah H, Sundararaju S, Dalil L, Abdelwahab AH, Salameh S, Ibrahim EB, Almaslmani MA, Tang P, Perez-Lopez A, Tsui CKM. 2023. Molecular characterization of *Candida auris* outbreak isolates in Qatar from patients with COVID-19 reveals the emergence of isolates resistant to three classes of antifungal drugs. Clin Microbiol Infect 29:1083.e1–1083.e7.

77. Spruijtenburg B, Ahmad S, Asadzadeh M, Alfouzan W, Al-Obaid I, Mokaddas E, Meijer EFJ, Meis JF, de Groot T. 2023. Whole genome sequencing analysis demonstrates therapy-induced echinocandin resistance in *Candida auris* isolates. Mycoses 10.1111/myc.13655.

78. Burrack LS, Todd RT, Soisangwan N, Wiederhold NP, Selmecki A. 2022. Genomic diversity across *Candida auris* clinical isolates shapes rapid development of antifungal resistance *in vitro* and *in vivo*. mBio 13:e0084222.

79. Jacobs SE, Jacobs JL, Dennis EK, Taimur S, Rana M, Patel D, Gitman M, Patel G, Schaefer S, Iyer K, Moon J, Adams V, Lerner P, Walsh TJ, Zhu Y, Anower MR, Vaidya MM, Chaturvedi S, Chaturvedi V. 2022. *Candida auris* pan-drug-resistant to four classes of antifungal agents. Antimicrob Agents Chemother 66:e0005322.

80. Gorzalski A, Ambrosio FJ, Massic L, Scribner MR, Siao DD, Hua C, Dykema P, Schneider E, Njoku C, Libuit K, Sevinsky JR, Van Hooser S, Pandori M, Hess D. 2023. The use of whole-genome sequencing and development of bioinformatics to monitor overlapping outbreaks of *Candida auris* in southern Nevada. Front Public Health 11:1198189.

81. Spruijtenburg B, Badali H, Abastabar M, Mirhendi H, Khodavaisy S, Sharifisooraki J, Taghizadeh Armaki M, De Groot T, Meis JF. 2022. Confirmation of fifth *Candida auris* clade by whole genome sequencing. Emerg Microbes Infect 11:2405–2411.

82. De Luca DG, Alexander DC, Dingle TC, Dufresne PJ, Hoang LM, Kus JV, Schwartz IS, Mulvey MR, Bharat A. 2022. Four genomic clades of *Candida auris* identified in Canada, 2012-2019. Med Mycol 60:myab079.

83. Tian S, Bing J, Chu Y, Chen J, Cheng S, Wang Q, Zhang J, Ma X, Zhou B, Liu L, Huang G, Shang H. 2021. Genomic epidemiology of *Candida auris* in a general hospital in Shenyang, China: a three-year surveillance study. Emerg Microbes Infect 10:1088–1096.

84. Salah H, Sundararaju S, Dalil L, Salameh S, Al-Wali W, Tang P, Ben Abid F, Tsui CKM. 2021. Genomic epidemiology of *Candida auris* in Qatar reveals hospital transmission dynamics and a south Asian origin. J Fungi 7:240.

85. Price TK, Mirasol R, Ward KW, Dayo AJ, Hilt EE, Chandrasekaran S, Garner OB, de St Maurice A, Yang S. 2021. Genomic characterizations of Clade III lineage of *Candida auris*, California, USA. Emerg Infect Dis 27:1223–1227.

86. Arora P, Singh P, Wang Y, Yadav A, Pawar K, Singh A, Padmavati G, Xu J, Chowdhary A. 2021. Environmental isolation of *Candida auris* from the coastal wetlands of Andaman Islands, India. mBio 12:e03181–20.

87. Biswas C, Wang Q, van Hal SJ, Eyre DW, Hudson B, Halliday CL, Mazsewska K, Kizny Gordon A, Lee A, Irinyi L, Heath CH, Chakrabarti A, Govender NP, Meyer W, Sintchenko V, Chen SC-A. 2020. Genetic heterogeneity of Australian *Candida auris* isolates: insights from a nonoutbreak setting using whole-genome sequencing. Open Forum Infect Dis 7:ofaa158.

88. Tan YE, Teo JQ-M, Rahman NBA, Ng OT, Kalisvar M, Tan AL, Koh TH, Ong RTH. 2019. *Candida auris* in Singapore: genomic epidemiology, antifungal drug resistance, and identification using the updated 8.01 VITEK2 system. Int J Antimicrob Agents 54:709– 715.

89. Sekizuka T, Iguchi S, Umeyama T, Inamine Y, Makimura K, Kuroda M, Miyazaki Y, Kikuchi K. 2019. Clade II *Candida auris* possess genomic structural variations related to an ancestral strain. PloS One 14:e0223433.

90. Vogelzang EH, Weersink AJL, van Mansfeld R, Chow NA, Meis JF, van Dijk K. 2019. The first two cases of *Candida auris* in the Netherlands. J Fungi Basel Switz 5:91.

91. Chow NA, de Groot T, Badali H, Abastabar M, Chiller TM, Meis JF. 2019. Potential fifth clade of *Candida auris*, Iran, 2018. Emerg Infect Dis 25:1780–1781.

92. Pchelin IM, Azarov DV, Churina MA, Ryabinin IA, Vibornova IV, Apalko SV, Kruglov AN, Sarana AM, Taraskina AE, Vasilyeva NV. 2020. Whole genome sequence of first *Candida auris* strain, isolated in Russia. Med Mycol 58:414–416.

93. Heath CH, Dyer JR, Pang S, Coombs GW, Gardam DJ. 2019. *Candida auris* sternal osteomyelitis in a man from Kenya visiting Australia, 2015. Emerg Infect Dis 25:192–194.

94. Chow NA, Gade L, Tsay SV, Forsberg K, Greenko JA, Southwick KL, Barrett PM, Kerins JL, Lockhart SR, Chiller TM, Litvintseva AP, US Candida auris Investigation Team. 2018. Multiple introductions and subsequent transmission of multidrug-resistant Candida auris in the USA: a molecular epidemiological survey. Lancet Infect Dis 18:1377–1384.

95. Rhodes J, Abdolrasouli A, Farrer RA, Cuomo CA, Aanensen DM, Armstrong-James D, Fisher MC, Schelenz S. 2018. Genomic epidemiology of the UK outbreak of the emerging human fungal pathogen *Candida auris*. Emerg Microbes Infect 7:43.

96. 2026. ncbi/sra-tools. C. NCBI - National Center for Biotechnology Information/NLM/NIH.

97. L. Hoyer L. 2023. Extraction of Yeast High-Molecular-Weight Genomic DNA v1 10.17504/protocols.io.rm7vzb1b4vx1/v1.

98. Boratyn GM, Thierry-Mieg J, Thierry-Mieg D, Busby B, Madden TL. 2019. Magic-BLAST, an accurate RNA-seq aligner for long and short reads. BMC Bioinformatics 20:405.

99. Cauldron NC, Shea T, Cuomo CA. 2024. Improved genome assembly of *Candida auris* strain *B8441* and annotation of *B11205*. Microbiol Resour Announc 0:e00512–24.

100. Altschul SF, Gish W, Miller W, Myers EW, Lipman DJ. 1990. Basic local alignment search tool. J Mol Biol 215:403–410.

101. Camacho C, Coulouris G, Avagyan V, Ma N, Papadopoulos J, Bealer K, Madden TL. 2009. BLAST+: architecture and applications. BMC Bioinformatics 10:421.

102. Price MN, Dehal PS, Arkin AP. 2010. FastTree 2 – approximately maximum-likelihood trees for large alignments. PLOS ONE 5:e9490.

103. Li H, Durbin R. 2009. Fast and accurate short read alignment with Burrows–Wheeler transform. Bioinformatics 25:1754–1760.

104. Li H, Handsaker B, Wysoker A, Fennell T, Ruan J, Homer N, Marth G, Abecasis G, Durbin R. 2009. The sequence alignment/map format and SAMtools. Bioinformatics 25:2078–2079.

105. McKenna A, Hanna M, Banks E, Sivachenko A, Cibulskis K, Kernytsky A, Garimella K, Altshuler D, Gabriel S, Daly M, DePristo MA. 2010. The Genome Analysis Toolkit: A MapReduce framework for analyzing next-generation DNA sequencing data. Genome Res 20:1297–1303.

106. Sephton-Clark P, Tenor JL, Toffaletti DL, Meyers N, Giamberardino C, Molloy SF, Palmucci JR, Chan A, Chikaonda T, Heyderman R, Hosseinipour M, Kalata N, Kanyama C, Kukacha C, Lupiya D, Mwandumba HC, Harrison T, Bicanic T, Perfect JR, Cuomo CA. 2022. Genomic variation across a clinical *Cryptococcus* population linked to disease outcome. mBio 13:e02626–22.

107. Cingolani P, Platts A, Wang LL, Coon M, Nguyen T, Wang L, Land SJ, Lu X, Ruden DM. 2012. A program for annotating and predicting the effects of single nucleotide polymorphisms, SnpEff: SNPs in the genome of *Drosophila melanogaster* strain w1118; iso-2; iso-3. Fly (Austin) 6:80–92.

108. Zhou X, Stephens M. 2012. Genome-wide efficient mixed-model analysis for association studies. Nat Genet 44:821–824.

109. Zhou X, Stephens M. 2014. Efficient multivariate linear mixed model algorithms for genome-wide association studies. Nat Methods 11:407–409.

110. Danecek P, Bonfield JK, Liddle J, Marshall J, Ohan V, Pollard MO, Whitwham A, Keane T, McCarthy SA, Davies RM, Li H. 2021. Twelve years of SAMtools and BCFtools. GigaScience 10:giab008.

111. Shimodaira H, Hasegawa M. 1999. Multiple comparisons of log-likelihoods with applications to phylogenetic inference. Mol Biol Evol 16:1114.

112. Sagulenko P, Puller V, Neher RA. 2018. TreeTime: maximum-likelihood phylodynamic analysis. Virus Evol 4:vex042.

113. Rice P, Longden I, Bleasby A. 2000. EMBOSS: The European Molecular Biology Open Software Suite. Trends Genet 16:276–277.

114. Robbins EHJ, Liu Y, Kelly S. RECUR: Identifying recurrent amino acid substitutions from multiple sequence alignments.

115. R Core Team. 2025. R: A language and environment for statistical computing. R Foundation for Statistical Computing, Vienna, Austria. https://www.R-project.org/.

116. Nei M, Gojobori T. 1986. Simple methods for estimating the numbers of synonymous and nonsynonymous nucleotide substitutions. Mol Biol Evol 3:418–426.

117. Schloissnig S, Arumugam M, Sunagawa S, Mitreva M, Tap J, Zhu A, Waller A, Mende DR, Kultima JR, Martin J, Kota K, Sunyaev SR, Weinstock GM, Bork P. 2013. Genomic variation landscape of the human gut microbiome. Nature 493:45–50.

118. dos Reis M. 2015. How to calculate the non-synonymous to synonymous rate ratio of protein-coding genes under the Fisher–Wright mutation–selection framework. Biol Lett 11:20141031.

119. Parto S, Lartillot N. 2018. Molecular adaptation in Rubisco: Discriminating between convergent evolution and positive selection using mechanistic and classical codon models. PLOS ONE 13:e0192697.

120. Gade L, Muñoz JF, Sheth M, Wagner D, Berkow EL, Forsberg K, Jackson BR, Ramos-Castro R, Escandón P, Dolande M, Ben-Ami R, Espinosa-Bode A, Caceres DH, Lockhart SR, Cuomo CA, Litvintseva AP. 2020. Understanding the emergence of multidrug-resistant *Candida*: Using whole-genome sequencing to describe the population structure of *Candida haemulonii* species complex. Front Genet 11.

121. Rivera-Colón AG, Rehmann CT, Kern AD. 2026. MKado: a toolkit for McDonald-Kreitman tests of natural selection. bioRxiv 10.64898/2026.03.02.709122.

122. Barker KS, Santana DJ, Zhang Q, Peters TL, Rybak JM, Morschhäuser J, Cuomo CA, Rogers PD. 2025. Mutations in *TAC1B* drive increased *CDR1* and *MDR1* expression and azole resistance in *Candida auris*. Antimicrob Agents Chemother 69:e00300–25.

123. Doorley LA, Meza-Perez V, Jones SJ, Rybak JM. 2025. A *Candidozyma* (*Candida*) *auris*-optimized Episomal Plasmid Induced Cas9-editing system reveals the direct impact of the S639F encoding *FKS1* mutation. J Infect Dis 2025.02.14.638356.

124. 2026. Benchling [Biology Software].

125. Lombardi L, Oliveira-Pacheco J, Butler G. 2019. Plasmid-based CRISPR-Cas9 gene editing in multiple *Candida* species. mSphere 4:10.1128/msphere.00125-19.

126. 2017. M27 Reference Method for Broth Dilution Antifungal Susceptibility Testing of Yeasts, 4th ed. Clinical and Laboratory Standards Institute (CLSI), Wayne, PA.

127. Rybak JM, Muñoz JF, Barker KS, Parker JE, Esquivel BD, Berkow EL, Lockhart SR, Gade L, Palmer GE, White TC, Kelly SL, Cuomo CA, Rogers PD. 2020. Mutations in *TAC1B*: a novel genetic determinant of clinical fluconazole resistance in *Candida auris*. mBio 11:10.1128/mbio.00365-20.

128. Rybak JM, Sharma C, Doorley LA, Barker KS, Palmer GE, Rogers PD. 2021. Delineation of the direct contribution of *Candida auris ERG11* mutations to clinical triazole resistance. Microbiol Spectr 9:e01585–21.

129. Liu F, Hu Z-D, Zhao X-M, Zhao W-N, Feng Z-X, Yurkov A, Alwasel S, Boekhout T, Bensch K, Hui F-L, Bai F-Y, Wang Q-M. 2024. Phylogenomic analysis of the *Candida auris-Candida haemuli* clade and related taxa in the Metschnikowiaceae, and proposal of thirteen new genera, fifty-five new combinations and nine new species. Persoonia - Mol Phylogeny Evol Fungi 52:22–43.

130. Ekenoğlu Merdan Y, Zıkşahna K. 2025. Beyond candidiasis: should infections caused by *Candidozyma auris* be renamed? J Clin Microbiol 0:e01530–25.

131. Zhang SX, de Hoog S, Denning DW, Ahmed SA, Alastruey-Izquierdo A, Arendrup MC, Borman A, Chen S, Chowdhary A, Colgrove RC, Cornely OA, Dufresne PJ, Filkins L, Gangneux J-P, Gené J, Groll AH, Guillot J, Haase G, Halliday C, Hawksworth DL, Hay R, Hoenigl M, Hubka V, Jagielski T, Kandemir H, Kidd SE, Kus JV, Kwon-Chung J, Lockhart SR, Meis JF, Mendoza L, Meyer W, Nguyen MH, Song Y, Sorrell TC, Stielow JB, Vilela R, Vitale RG, Wengenack NL, White PL, Ostroski-Zeichner L, Walsh TJ, members of the ISHAM/ECMM/FDLC Working Group Nomenclature of Clinical Fungi. 2025. Reaffirming the importance of nomenclature stability for Candida auris and its associated disease of candidiasis. J Clin Microbiol 0:e01550–25.

132. Arnaud MB, Costanzo MC, Skrzypek MS, Binkley G, Lane C, Miyasato SR, Sherlock G. 2005. The *Candida* Genome Database (CGD), a community resource for *Candida albicans* gene and protein information. Nucleic Acids Res 33:D358–D363.

133. Cherry JM, Hong EL, Amundsen C, Balakrishnan R, Binkley G, Chan ET, Christie KR, Costanzo MC, Dwight SS, Engel SR, Fisk DG, Hirschman JE, Hitz BC, Karra K, Krieger CJ, Miyasato SR, Nash RS, Park J, Skrzypek MS, Simison M, Weng S, Wong ED. 2012. *Saccharomyces* Genome Database: the genomics resource of budding yeast. Nucleic Acids Res 40:D700–D705.

134. Lew-Smith J, Binkley J, Sherlock G. 2025. The *Candida* Genome Database: annotation and visualization updates. Genetics 229:iyaf001.

135. Staab JF, Neofytos D, Rhee P, Jiménez-Ortigosa C, Zhang SX, Perlin DS, Marr KA. 2014. Target enzyme mutations confer differential echinocandin susceptibilities in *Candida kefyr*. Antimicrob Agents Chemother 58:5421–5427.

136. Cowen LE, Sanglard D, Howard SJ, Rogers PD, Perlin DS. 2015. Mechanisms of antifungal drug resistance. Cold Spring Harb Perspect Med 5:a019752.

137. Prigent G, Aït-Ammar N, Levesque E, Fekkar A, Costa J-M, El Anbassi S, Foulet F, Duvoux C, Merle J-C, Dannaoui E, Botterel F. 2017. Echinocandin resistance in *Candida* species isolates from liver transplant recipients. Antimicrob Agents Chemother 61:10.1128/aac.01229-16.

138. Hirayama T, Miyazaki T, Yamagishi Y, Mikamo H, Ueda T, Nakajima K, Takesue Y, Higashi Y, Yamamoto Y, Kimura M, Araoka H, Taniguchi S, Fukuda Y, Matsuo Y, Furutani A, Yamashita K, Takazono T, Saijo T, Shimamura S, Yamamoto K, Imamura Y, Izumikawa K, Yanagihara K, Kohno S, Mukae H. 2018. Clinical and microbiological characteristics of *Candida guilliermondii* and *Candida fermentati*. Antimicrob Agents Chemother 62:10.1128/aac.02528-17.

139. Silva LN, Ramos LS, Oliveira SSC, Magalhães LB, Cypriano J, Abreu F, Macedo AJ, Branquinha MH, Santos ALS. 2023. Development of echinocandin resistance in *Candida haemulonii*: an emergent, widespread, and opportunistic fungal pathogen. J Fungi 9:859.

140. Mio T, Adachi-Shimizu M, Tachibana Y, Tabuchi H, Inoue SB, Yabe T, Yamada-Okabe T, Arisawa M, Watanabe T, Yamada-Okabe H. 1997. Cloning of the *Candida albicans* homolog of *Saccharomyces cerevisiae GSC1*/*FKS1* and its involvement in beta-1,3-glucan synthesis. J Bacteriol 179:4096–4105.

141. Basenko EY, Pulman JA, Shanmugasundram A, Harb OS, Crouch K, Starns D, Warrenfeltz S, Aurrecoechea C, Stoeckert CJ, Kissinger JC, Roos DS, Hertz-Fowler C. 2018. FungiDB: an integrated bioinformatic resource for fungi and oomycetes. J Fungi 4:39.

142. Emms DM, Kelly S. 2019. OrthoFinder: phylogenetic orthology inference for comparative genomics. Genome Biol 20:238.

143. Emms DM, Kelly S. 2022. SHOOT: phylogenetic gene search and ortholog inference. Genome Biol 23:85.

144. Buchfink B, Xie C, Huson DH. 2015. Fast and sensitive protein alignment using DIAMOND. 1. Nat Methods 12:59–60.

145. Zheng J, Ge Q, Yan Y, Zhang X, Huang L, Yin Y. 2023. dbCAN3: automated carbohydrate-active enzyme and substrate annotation. Nucleic Acids Res 51:W115–W121.

146. Katoh K, Standley DM. 2013. MAFFT multiple sequence alignment software version 7: improvements in performance and usability. Mol Biol Evol 30:772–780.

147. Steenwyk JL, Iii TJB, Li Y, Shen X-X, Rokas A. 2020. ClipKIT: A multiple sequence alignment trimming software for accurate phylogenomic inference. PLOS Biol 18:e3001007.

148. Suyama M, Torrents D, Bork P. 2006. PAL2NAL: robust conversion of protein sequence alignments into the corresponding codon alignments. Nucleic Acids Res 34:W609–W612.

149. Kozlov AM, Darriba D, Flouri T, Morel B, Stamatakis A. 2019. RAxML-NG: a fast, scalable and user-friendly tool for maximum likelihood phylogenetic inference. Bioinformatics 35:4453–4455.

150. Kozlov AM, Stamatakis A. 2020. Using RAxML-NG in Practice, p. 1.3:1-1.3:25. In Phylogenetics in the Genomic Era. Authors open access book.

151. Darriba D, Posada D, Kozlov AM, Stamatakis A, Morel B, Flouri T. 2020. ModelTest-NG: a new and scalable tool for the selection of DNA and protein evolutionary models. Mol Biol Evol 37:291–294.

152. Yang Z. 2007. PAML 4: Phylogenetic Analysis by Maximum Likelihood. Mol Biol Evol 24:1586–1591.

153. Álvarez-Carretero S, Kapli P, Yang Z. 2023. Beginner’s guide on the use of PAML to detect positive selection. Mol Biol Evol 40:msad041.

154. Pettersen EF, Goddard TD, Huang CC, Meng EC, Couch GS, Croll TI, Morris JH, Ferrin TE. 2021. UCSF ChimeraX: Structure visualization for researchers, educators, and developers. Protein Sci 30:70–82.

155. Abramson J, Adler J, Dunger J, Evans R, Green T, Pritzel A, Ronneberger O, Willmore L, Ballard AJ, Bambrick J, Bodenstein SW, Evans DA, Hung C-C, O’Neill M, Reiman D, Tunyasuvunakool K, Wu Z, Žemgulytė A, Arvaniti E, Beattie C, Bertolli O, Bridgland A, Cherepanov A, Congreve M, Cowen-Rivers AI, Cowie A, Figurnov M, Fuchs FB, Gladman H, Jain R, Khan YA, Low CMR, Perlin K, Potapenko A, Savy P, Singh S, Stecula A, Thillaisundaram A, Tong C, Yakneen S, Zhong ED, Zielinski M, Žídek A, Bapst V, Kohli P, Jaderberg M, Hassabis D, Jumper JM. 2024. Accurate structure prediction of biomolecular interactions with AlphaFold 3. Nature 630:493–500.

