## Supplementary Material for "Parallel evolution under constraint shapes echinocandin resistance in *Candida auris*"

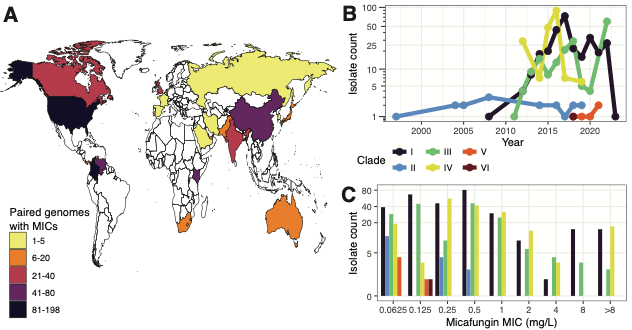

**Supplementary Fig. 1. Collections of 681 *C. auris* isolates with genomic and drug susceptibility data available span the globe and historic outbreaks. A)** World map colored by the number of isolates from each country that are included in the study. **B)** Isolates were collected between 1997-2023, though there is only one isolate from each of 1997 and 2023, and none spanning 1998-2003. The absence of more contemporaneous samples is likely an artifact of delay between sample collection and original publication, compounded by the delay between when our search was performed and article submission. **C)** Micafungin MIC distribution for included isolates.

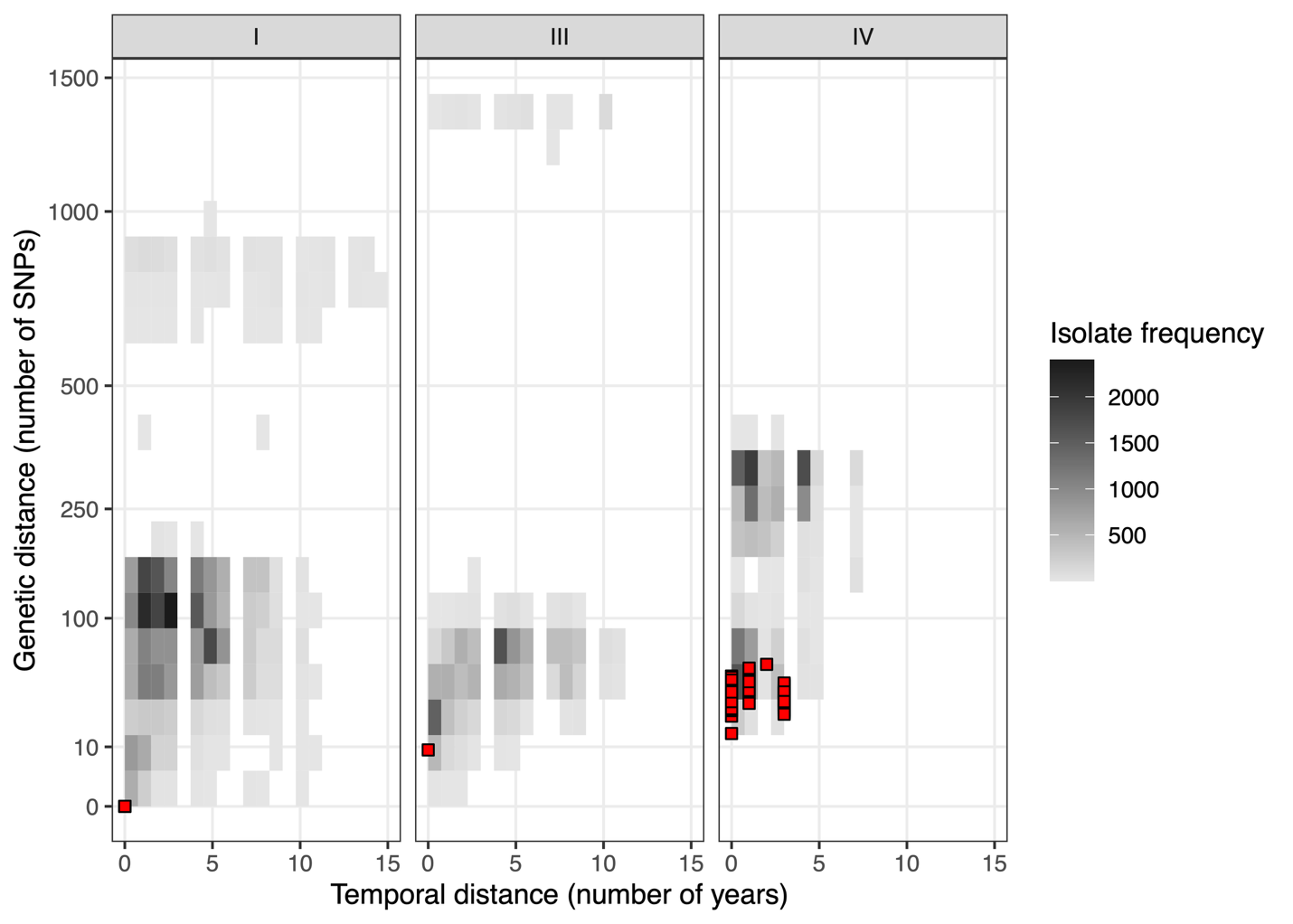

**Supplementary Fig. 2. Candidate transmission clusters of *FKS1*-driven micafungin resistance are a low number of SNPs apart against the genetic background of their clade.** Red points indicate distances within predicted transmission clusters.

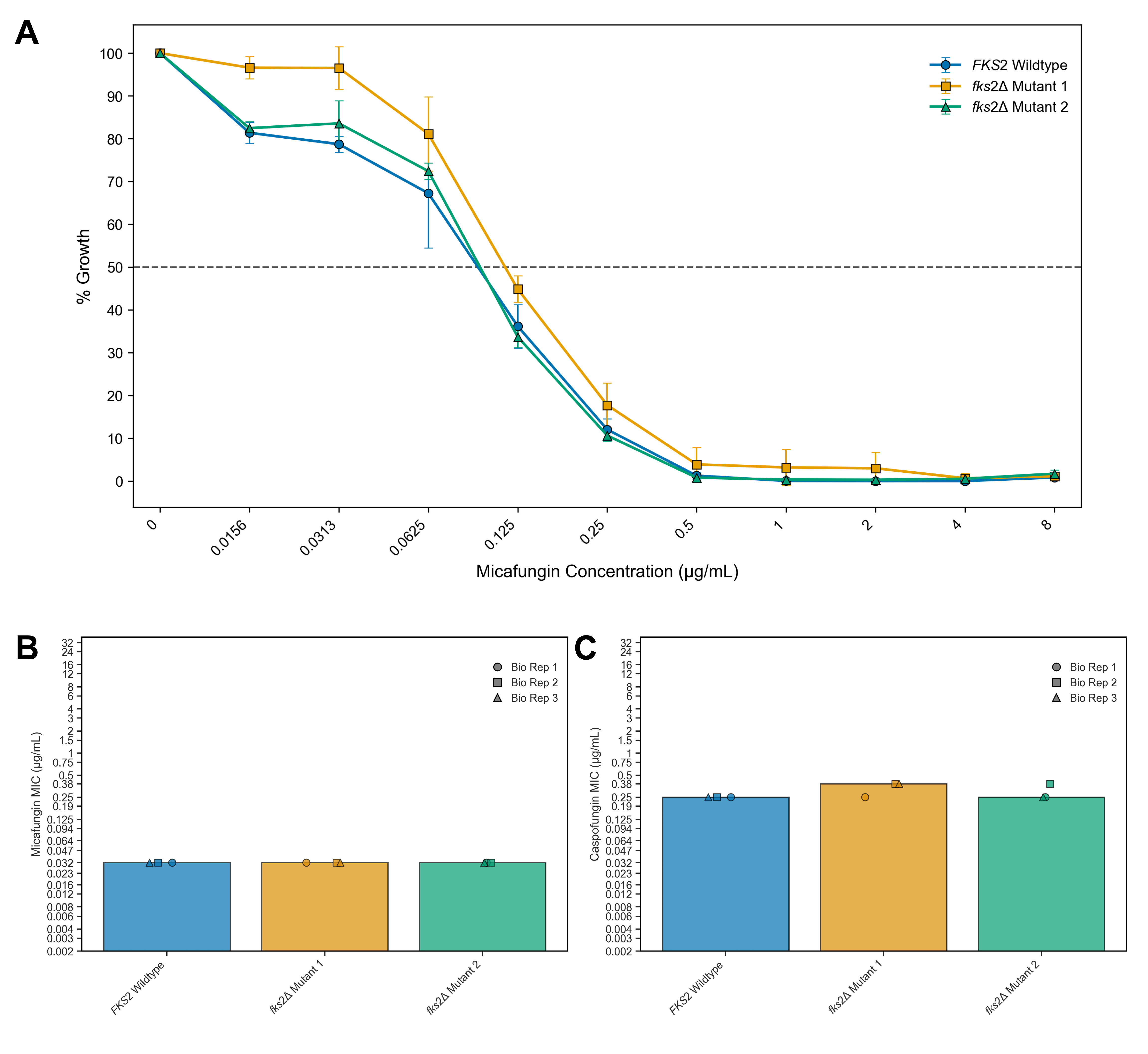

**Supplementary Fig. 3. Echinocandin antifungal susceptibility testing of *C. auris* 1cF (*FKS2*) and two independent disruption mutant (*fks2*Δ) strains.** A) Micafungin broth microdilution results shown as percent growth relative to the corresponding DMSO control after 24 hours incubation with increasing drug concentrations. Testing was performed in biological triplicate and technical duplicate. Each data point represents the mean of the three biological replicates ± standard deviation. The horizontal dotted line represents 50% growth inhibition. Gradient diffusion strip results for B) micafungin, and C) caspofungin shown as modal MICs across three biological replicates.

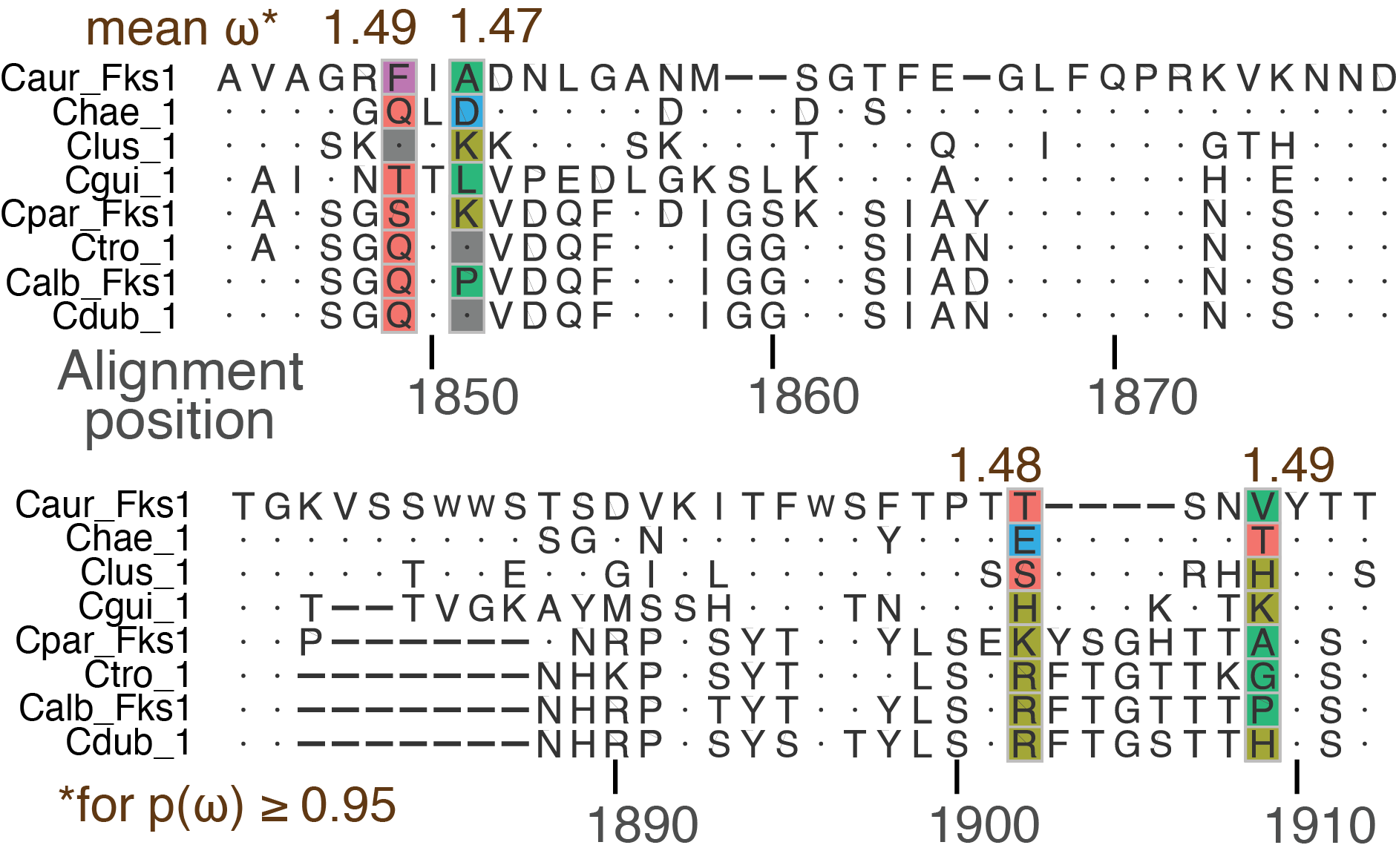

**Supplementary Fig. 4.** **Multiple sequence alignment of the Fks1 region harboring positively selected sites among species in the CTG clade**. The estimated selection coefficient ω is shown near the positions with a coefficient significantly different from the theoretical expectation of a neutrally evolving site (BEB; p(ω>1) ≥ 0.95). Taxa are abbreviated: *C. auris* (Cau), *C. lusitaniae* (Clus), *C. haemulonii* (Chae), *C. albicans* (Calb), *C. dubliniensis* (Cdub), *C. tropicalis* (Ctrop), *C. guillermondii* (Cgui), *C. parapsilosis* (Cpar), *C. krusei* (Ckru), *C. kefyr* (Ckef), *C. glabrata* (Cgla) and *S. cerevisiae* (Scer).

**Supplementary Table 1. Distribution of micafungin MICs (µg/mL) among isolates analyzed in either GWAS performed.**

| **Micafungin MIC**** | **Susceptibility index*** | **All isolates**† | **Fks1 hotspot variants excluded** |
| --- | --- | --- | --- |
| <= 0.125 | <= 0.5 | 323 | 330 |
| 0.25 | 1 | 170 | 170 |
| 0.5 | 2 | 87 | 87 |
| 1 | 3 | 29 | 27 |
| 2 | 4 | 8 | 7 |
| 4 | 5 | **17** | **1** |
| 8 | 6 | **29** | **0** |
| > 8 | 7 | **6** | **1** |

** Data obtained from publications or shared by the CDC.

* The susceptibility index is calculated as log2 dilution of MIC value larger than 0.25 µg/mL. MICs less than 0.25 µg/mL were calculated as a linear proportion of this value.

† Isolate counts in bold are resistant relative to the CDC’s tentative breakpoint (4.0 µg/mL).

**Supplementary Table 2. The number of Fks1 mutational events predicted to occur in the ancestry of each clade and MICs of isolates harboring those mutations.**

| **Fks1 allele** | **Hotspot** | **Clade** | **Mut. events** | **# Derived mutations*** | **# Ancestral mutations**† | **# Total** | **Modal MIC** | **Median MIC** |
| --- | --- | --- | --- | --- | --- | --- | --- | --- |
| S639F | 1 | I | 9 | 7 | 4 | 11 | 8 | 8 |
| S639Y | 1 |  | 8 | 5 | 6 | 11 | 4 | 4 |
| S639P | 1 |  | 2 | 1 | 3 | 4 | 4 | 4 |
| D642Y | 1 |  | 1 | 1 | 0 | 1 | 0.25 | 0.25 |
| L1357F | 2 |  | 1 | 1 | 0 | 1 | 0.06 | 0.06 |
| L638F | 1 |  | 1 | 1 | 0 | 1 | 0.5 | 0.5 |
| M690V | 3 |  | 1 | 1 | 0 | 1 | 8 | 8 |
| R641S | 1 |  | 1 | 1 | 0 | 1 | 2 | 2 |
| F635C | 1 |  | 1 | 1 | 0 | 1 | 8 | 8 |
| F635del,F1367L | 1, Non-HS |  | 1 | 1 | 0 | 1 | 8 | 8 |
| *WT* | *-* |  | - | - | - | 259 | 0.25 | 0.125 |
| L1572I | Non-HS | II | 1 | 0 | 17 | 17 | 0.06 | 0.06 |
| S639F | 1 | III | 3 | 3 | 0 | 3 | 8 | 8 |
| S639Y | 1 |  | 1 | 1 | 0 | 1 | 4 | 4 |
| F635dup | 1 |  | 1 | 0 | 2 | 2 | 4,0.03 | 2.015 |
| *WT* | *-* |  | - | - | - | 162 | 0.25 | 0.1875 |
| S639P | 1 | IV | 2 | 0 | 18 | 18 | 8 | 8 |
| M1267I | Non-HS |  | 1 | 0 | 6 | 6 | 1 | 1 |
| S639F,L1357F | 1, 2 |  | 1 | 1 | 0 | 1 | 8 | 8 |
| *WT* | *-* |  | - | - | - | 161 | 0.125 | 0.25 |
| I1817V | Non-HS | V | 1 | 0 | 5 | 5 | 0.031 | 0.031 |
| *WT* | *-* | VI | - | - | - | 1 | 0.12 | 0.12 |

* Mutations at terminal phylogenetic nodes were considered to have been acquired independently

† Mutations at internal phylogenetic nodes were considered ancestral to clusters

**Supplementary Table 3. Missense mutations in *FKS1* and *FKS2* among *C. auris* coding DNA sequences (n=22,388);** **attached as Supplementary File 2.**

**Supplementary Table 4. *FKS* genes of *C. auris* and *S. cerevisiae* cluster into three hierarchical orthologous groups (HOGs).**

| **Gene ID** | **Gene Symbol** | **HOG** |
| --- | --- | --- |
| **B9J08_000964** | **Caur_FKS1** | 1 |
| **C1_02420C_A** | **Calb_GSC1** | 1 |
| C5L36_0E04550 | Ckru_1 | 1 |
| **CAGL0G01034g** | **Cgla_FKS1** | 1 |
| **CAGL0K04037g** | **Cgla_FKS2** | 1 |
| CD36_02270 | Cdub_1 | 1 |
| **CPAR2_106400** | **Cpar_FKS1** | 1 |
| CTMYA2_028570 | Ctro_1 | 1 |
| CXQ85_001170 | Chae_1 | 1 |
| FOB63_001684 | Clus_1 | 1 |
| KLMA_50197 | Ckef_1 | 1 |
| PGUG_01168 | Cgui_1 | 1 |
| **YGR032W** | **Scer_GSC2** | 1 |
| **YLR342W** | **Scer_FKS1** | 1 |
| **B9J08_001020** | **Caur_FKS2** | 2 |
| CD36_25900 | Cdub_2 | 2 |
| **CPAR2_804030** | **Cpar_FKS2** | 2 |
| **CR_00850C_A** | **Calb_GSL2** | 2 |
| CTMYA2_042050 | Ctro_2 | 2 |
| CXQ85_001220 | Chae_2 | 2 |
| KLMA_30497 | Ckef_2 | 2 |
| PGUG_00803 | Cgui_2 | 2 |
| C5L36_0B08470 | Ckru_3 | 3 |
| **CAGL0M13827g** | **Cgla_FKS3** | 3 |
| KLMA_80197 | Ckef_3 | 3 |
| **YMR306W** | **Scer_FKS3** | 3 |

**Supplementary Table 5. Branch selection tests on *FKS1* and *FKS2* among Saccharomycetes.**

| **Gene** | **Model** | **dS tree length** | ***C. auris* dN/dS** | **lnL** | **np** | **df vs M0** | **LRT vs M0** | **P-value vs M0** |
| --- | --- | --- | --- | --- | --- | --- | --- | --- |
| CTG - excluding Cgui and Clus | |  |  |  |  |  |  |  |
| *FKS1* | M0 | 3.52 | 0.045 | -18070.2 | 12 |  |  |  |
|  | M1 | 3.13 | 0.025 | -18040.3 | 21 | 21 | -36081 | 1.6E-09 |
| *FKS2* | M0 | 30.45 | 0.019 | -24449.5 | 12 |  |  |  |
|  | M1 | 87.81 | 0.046 | -24442.1 | 21 | 21 | -48884 | 9.7E-02 |
| CTG |  |  |  |  |  |  |  |  |
| *FKS1* | M0 | 4.45 | 0.060 | -23637.3 | 16 |  |  |  |
|  | M1 | 4.21 | 0.016 | -23555.0 | 29 | 29 | -47110 | 2.4E-28 |
| Saccharomycetes |  |  |  |  |  |  |  |  |
| *FKS1* | M0 | 510.02 | 0.039 | -87561.9 | 27 |  |  |  |
|  | M1 | 567.01 | 0.033 | -87532.3 | 53 | 26 | -175065 | 2.2E-04 |

**Supplementary Table 6. Site selection tests on *FKS1* among the CTG clade in *Candida*.**

| **Model** | **lnL** | **n.p.** | **Model comparison** | **d.f.** | **LRT** | **p-value** |
| --- | --- | --- | --- | --- | --- | --- |
| M0 | -23637.298 | 16 |  |  |  |  |
| M1a | -23229.301 | 17 | M0 vs. M1a (one-ratio vs. nearly-neutral) | 1 | 816.00 | 0.000 |
| M2a | -23229.302 | 19 | M1a vs. M2a (nearly-neutral vs. positive selection) | 2 | 0.00 | - |
| M7 | -23077.759 | 17 |  |  |  |  |
| M8a | -23072.112 | 18 | M8a vs. M8 (beta&w with w=1) | 1 | 5.56 | 0.018 |
| M8 | -23066.556 | 19 | M7 vs. M8 (beta vs. beta&w with w>1) | 2 | 22.41 | < 0.001 |

**Supplementary Table 7.** **Oligonucleotides used in this study for genetic transformation of *C. auris*.**

| **Oligonucleotide name** | **Oligonucleotide sequence (5’–3’)** |
| --- | --- |
| FKS2dis27 gRNA-TOP | CCAAGAAGTTATCACCCCCCAAG |
| FKS2dis27 gRNA-BOT | AACCTTGGGGGGTGATAACTTCT |
| FKS2dis27RT (gBlock repair template) | TCAGCATCTTTCGTGATGTCGGTTTTGACAGACAGCTTCTCTTCAGCGTAGCACCGAAAGAAGCGATCAGGCCCGTGGCCCATTCCCTCGACGCCAGCCCAATGGGACGAGTCGAGGTACGGGGTTCGAAACGCCGACAATGGCGCCACGGCCGTCTGGAGCCCAGTCCGGGCAACACCTTTCAGGTCACTGTTGGTATCGCCCAAATCAAACTGGTCTGTCAACGCGTCACCGTCAATAGGTGGATCTGAAATGTTAGAACAGAGAGGGGCTGAGAAACTACGCGTTTGGGTTTGGGGTCATGTTCATACTGGAAGATAGTGGTTGCTTTCTCTTGGGGGGTCTAACTATCTATACGCCCATGATCACAAAGATTTGAAGAAAAGAAACTGGAAGAAAATTGAATTCAGGAGTAGCGGATTTCGTGGGCTTTATATACCCCTGCGCCGCACACCCAGATAGGTTGCATAGTGAAAAGTGTAGTCCTTTGTTTAGCCTCGAAAACAGGAGAAACAGCACGAAGCTTTTCGATAACAAAGCACTTGAATTCTTCGGAACCCCGTCGGTTGGCAAACTGATGGCTACAAAATGAATAAGTGATTTAGTCCCGAGAAACCTTCATCAATAGTGGAAGCTCCACGCGTCTGCTGTCAGTACGATTC |
| FKS2dis27RT AMP-F | TCAGCATCTTTCGTGATGTCGG |
| FKS2dis27RT AMP-R | GAATCGTACTGACAGCAGACGC |
| FKS2dis27RT CHECK-F | ATCCTCCTGGAAGCCAAACACC |
| FKS2dis27RT CHECK-R | GGTTTCTCGGAAGAAAGTGGTGG |
